# Effectiveness of antifungal treatments during chytridiomycosis epizootics in populations of an endangered frog

**DOI:** 10.1101/2021.06.13.448228

**Authors:** Roland A. Knapp, Maxwell B. Joseph, Thomas C. Smith, Ericka E. Hegeman, Vance T. Vredenburg, James E. Erdman, Daniel M. Boiano, Andrea J. Jani, Cheryl J. Briggs

**Author notes:** Note: Author contributions are described in **Supporting Information**.

## Abstract

The recently-emerged amphibian chytrid fungus *Batrachochytrium dendrobatidis* (Bd) has had an unprecedented impact on global amphibian populations, and highlights the urgent need to develop effective mitigation strategies against this pathogen. We conducted field antifungal treatment experiments in populations of the endangered mountain yellow-legged frog during or immediately after Bd-caused mass die-off events. The objective of the treatments was to reduce Bd infection intensity (“load”) and in doing so alter frog-Bd dynamics and increase the probability of frog population persistence despite ongoing Bd infection. Experiments included treatment of early life stages (tadpoles and subadults) with the antifungal drug itraconazole, treatment of adults with itraconazole, and augmentation of the skin microbiome of subadults with *Janthinobacterium lividum*, a commensal bacterium with antifungal properties. All itraconazole treatments caused immediate reductions in Bd load, and produced longer-term effects that differed between life stages. In experiments focused on early life stages, Bd load was reduced in the two months immediately following treatment and was associated with increased survival of subadults. However, Bd load and frog survival returned to pre-treatment levels in less than one year, and treatment had no effect on population persistence. In adults, treatment reduced Bd load and increased frog survival over the three-year post-treatment period, consistent with frogs having developed an effective adaptive immune response against Bd. Despite this protracted period of reduced impacts of Bd on adults, recruitment of new individuals into the adult population was limited and the population eventually declined to near-extirpation. In the microbiome augmentation experiment, bathing frogs in a *J. lividum* solution after Bd load reduction with itraconazole increased concentrations of this bacterium on frogs, but concentrations declined to baseline levels within one month and did not have a protective effect against Bd infection. Collectively, these results suggest that Bd mitigation efforts focused on frog populations that have recently declined due to Bd emergence are ineffective in causing long-term changes in frog-Bd dynamics and increasing population persistence, due largely to the inability of early life stages to mount an effective immune response against Bd and resulting high susceptibility. This results in repeated recruitment failure and a low probability of population persistence in the face of ongoing Bd infection.

## Introduction

Emerging infectious diseases are increasingly common in wildlife, often due to anthropogenic changes in the ecology of the host or pathogen (Daszak et al. 2000, Cunningham et al. 2017). Impacts of disease on wildlife can be severe, including long-term population decline and even extinction, with far-reaching effects on species, communities, and ecosystems (Ostfeld et al. 2008, Scheele et al. 2019). Diseases of wildlife can also spill over to humans and domestic animals (Alexander et al. 2018). Collectively, these impacts of emerging wildlife diseases have significant consequences to global biodiversity and public health (Daszak et al. 2000). As such, the ability to control diseases in wildlife is critically important, but disease management is often difficult because wildlife diseases are relatively poorly described, many fewer intervention measures (e.g., vaccines) are available than for humans, and free ranging wildlife are inherently difficult to study and treat (Joseph et al. 2013). As a result, available management strategies are mostly insufficient to mitigate the destructive effects of disease on wildlife.

The amphibian disease chytridiomycosis is caused by the chytrid fungus *Batrachochytrium dendrobatidis* (“Bd”). This recently discovered pathogen (Berger et al. 1998, Longcore et al. 1999) is thought to have originated in Asia (O’Hanlon et al. 2018) and spread globally via human commerce (Schloegel et al. 2009). Bd is highly pathogenic to a wide range of amphibian taxa and, by one estimate, has caused the severe decline or extinction of at least 500 amphibian species (Scheele et al. 2019), with many more predicted to be at risk (Rödder et al. 2009). In an effort to reduce the impact of chytridiomycosis, several mitigation measures have been suggested as means to increase the fraction of frogs surviving chytridiomycosis outbreaks, including treating frogs with antifungal agents (using drugs or augmentation of the skin microbiome with probiotics), treating the environment with antifungals to reduce the pool of infectious zoospores, and reducing host population density (Woodhams et al. 2011, Garner et al. 2016). However, recent mathematical modeling suggests that none of these three types of mitigation measures are likely to be universally effective at preventing chytridiomycosis-induced population extirpation, but treating frogs had the greatest likelihood of a beneficial outcome (Drawert et al. 2017). Relatively few field tests of these treatment strategies have been conducted to date, and results from these trials indicate limited effectiveness in promoting host population persistence (Woodhams et al. 2012, Garner et al. 2016). As such, despite years of research on Bd-host dynamics and possible mitigation measures, field-tested methods to prevent ongoing Bd-caused amphibian declines and extinctions are still lacking.

Mountain yellow-legged (“MYL”) frogs are emblematic of global amphibian declines, including those caused by Bd. The MYL frog is a complex of two closely-related species, *Rana muscosa* and *Rana sierrae*, endemic to the mountains of California and adjacent Nevada, USA (Vredenburg et al. 2007). During the past century, MYL frogs have disappeared from more than 90% of their historical localities (Vredenburg et al. 2007) and are listed as “endangered” under the U.S. Endangered Species Act (U.S. Fish and Wildlife Service 2002, 2014). In the Sierra Nevada portion of their range, the primary causes of decline are the introduction of nonnative fish into naturally fishless water bodies and, more recently, the spread of Bd (Knapp and Matthews 2000, Vredenburg et al. 2010). MYL frogs are highly susceptible to chytridiomycosis. Arrival of Bd in a naive population typically results in rapid increases in Bd prevalence and infection intensity (“load”), and subsequent mass frog die-offs (Vredenburg et al. 2010). Such epizootics (epizootic = outbreak of disease in a wildlife population) generally lead to extirpation of the affected frog population, and hundreds of such extirpations have occurred in the past several decades as Bd spread across the Sierra Nevada (e.g., Rachowicz et al. 2006, Vredenburg et al. 2010). Examples of affected populations transitioning to an enzootic state, in which host populations coexist with Bd in a relatively stable dynamic, are rare (Briggs et al. 2010).

Empirical and modeling results from MYL frog populations indicate the primary role of Bd load in driving epizootics, and provide insights into the factors that might produce epizootic versus enzootic dynamics (Briggs et al. 2010, Vredenburg et al. 2010). Models developed for this system follow the number of infective Bd zoospores in a “zoospore pool” (i.e, a lake containing a population frogs) and the load on each frog. The growth rate of Bd is assumed to be an increasing function of host density, and frog mortality occurs when Bd load exceeds a threshold value (Briggs et al. 2010). Model results demonstrate that following Bd invasion into a naive host population, host extinction versus persistence can result solely from density-dependent host-pathogen dynamics. This suggests that suppression of Bd loads (e.g., by reducing frog density or treating frogs with antifungal agents), could increase frog survivorship and the likelihood of long-term population persistence in an enzootic state. Given the typical abundance of early life stages in naive MYL frog populations and their importance in driving frog-Bd dynamics (Briggs et al. 2010), treatments focused on early life stages could be particularly influential.

Although density-dependent frog-Bd dynamics alone can produce both epizootic and enzootic states, an adaptive immune response by frogs against Bd may increase the likelihood of an enzootic outcome (Woodhams et al. 2011). In contrast to the relatively low immunocompetence of early life stages (Rollins-Smith 1998, Grogan et al. 2018b), adults of some species, including MYL frogs, can develop adaptive immune defenses that may be at least partially protective against Bd (McMahon et al. 2014, Ellison et al. 2015, Grogan et al. 2018a). Antifungal treatments conducted during epizootics could slow the growth of Bd and allow the full development of adaptive immunity, which in turn could increase adult survival and population persistence (Woodhams et al. 2011). Treatment of frogs using antifungal drugs is typically conducted as a short-term pulse perturbation, and the ability of such short-term actions to cause long-lasting changes in frog-Bd dynamics is uncertain. In contrast, a press perturbation that, by definition, is sustained over a longer time period, may have a higher probability of producing long-lasting outcomes. One such press perturbation is the manipulation of the amphibian skin microbiome to shift the ambient microbial community to an alternative stable configuration that has stronger antifungal properties (Bletz et al. 2013, Woodhams et al. 2014). The feasibility of such a manipulation is suggested by laboratory experiments in which augmentation of the skin microbiome with antifungal bacteria altered frog-Bd dynamics and increased frog survival (Harris et al. 2009, Kueneman et al. 2016, but see also Becker et al. 2011).

During 2009-2018, we conducted six field trials of antifungal treatments applied to *R. sierrae* populations during or soon after epizootics, with a goal of reducing Bd load and increasing frog survival and population persistence. Based on multiple years of skin swab collection (e.g., Vredenburg et al. 2010), all study populations were Bd-naive in the years prior to the epizootic and subsequent treatment. Trials included (1) two treatments of early life stages (tadpoles and recently metamorphosed subadults) with the antifungal drug itraconazole (Garner et al. 2009), (2) three treatments of adults with itraconazole, and (3) one manipulation of the frog skin microbiome that involved exposing subadult frogs to *Janthinobacterium lividum*, a symbiotic bacterium on amphibian skin that has antifungal properties (Brucker et al. 2008).

All six antifungal treatments changed frog-Bd dynamics, and most increased frog survival. However, they failed to accomplish the ultimate objective of increasing the probability of frog population persistence, and all populations declined to extirpation or near-extirpation during the study periods. Nevertheless, the detailed results from multiple treatments across different life stages is unusual, and collectively allow important insights into the repeatability of treatment outcomes and the reasons for the failure of treatments to influence population persistence. These insights are essential for the future development of mitigation measures that accomplish this important objective in the face of one of the most devastating wildlife diseases in recorded history (Scheele et al. 2019).

## Methods

This section is divided into general methods that apply to all or most of the treatments, followed by a description of specific methods related to each treatment.

### General methods

#### Visual encounter surveys

We counted *R. sierrae* of all life stages (adults: *≥* 40 mm snout-vent length (SVL); subadults: < 40 mm; tadpoles) using diurnal visual encounter surveys (VES) of the entire water body shoreline and the first 100 m of inlet and outlet streams. Counts are highly repeatable (Knapp and Matthews 2000), but underestimate the number of animals present.

#### Capture-mark-recapture surveys

To allow estimation of adult survival, we used capture-mark-recapture (CMR) surveys (Joseph and Knapp 2018). During each summer, we re-visited the study lakes one to three times (i.e., primary periods), and during each primary period all frog populations were generally surveyed on either one day or on three consecutive days. During each daily survey, any adult frogs observed were captured, identified via their passive integrated transponder (PIT) tag (or tagged if untagged), and released. When captured for the first time during a primary period, frogs were also swabbed (see next section for details), measured, and weighed.

#### Quantifying Bd load using skin swabs

We quantified Bd load using standard swabbing and quantitative PCR methods (Boyle et al. 2004, Hyatt et al. 2007, see Vredenburg et al. 2010 for swabbing methods specific to MYL frogs). We defined Bd load as the number of ITS1 copies per swab (see Joseph and Knapp 2018 for details). For reference to figures provided in the Results, in post-metamorphic *R. sierrae*, Bd loads indicative of severe chytridiomycosis are *≥* 600,000 ITS copies (= 5.8 ITS copies on a log_10_ scale; Vredenburg et al. 2010, Joseph and Knapp 2018).

#### Itraconazole treatment

In each of the antifungal treatments, we captured adults, subadults, or tadpoles (depending on the treatment) from the study lakes using hand-held nets. Immediately following capture, we collected a skin swab sample from all animals or a subset (depending on the treatment) to describe Bd load. In addition, adults and subadults were measured and weighed, and adults were tagged using 8 mm PIT tags inserted under the dorsal skin via a small incision.

We held animals assigned to the “treated” group in large mesh pens (2 m x 2 m x 0.75 m) for the duration of the multi-day treatment period. Pens were anchored in the littoral zone of the study lakes, and contained shallow water and shoreline habitats for basking and deeper water habitat (up to 0.7 m) that frogs and tadpoles use at night (Figure S1). After swabbing, animals assigned to the untreated “control” group were held in pens only 3-24 hr and then released back into the lake.

Although it would have been ideal to hold animals from both the treated and control groups in pens for the duration of the treatment period, doing so could have produced spurious and misleading results. Bd transmission is expected to increase with frog density (Rachowicz and Briggs 2007), and holding untreated control animals in pens at relatively high density could therefore have increased their Bd loads and reduced survival more than would be expected for animals in the treated group that were given daily antifungal baths. This would have biased the outcome toward lower survival of control animals compared to treated animals even if the antifungal treatment itself had no effect on survival. Assuming that holding animals in pens for several days has some negative effect (due to increased Bd transmission even in treated frogs, and lack of feeding opportunities), if our study design caused biases they should be conservative, i.e., reducing the survival of treated animals relative to control animals.

To conduct the antifungal treatments, on each day during the multi-day treatment period we transferred all animals in the treated group from pens to small plastic tubs that contained a dilute solution (1.5 mg L^-1^; Garner et al. 2009) of the antifungal drug itraconazole (trade name = Sporonox). The volume of itraconazole solution varied between 2 and 5 L and allowed all life stages to submerge fully. We treated frogs in batches of approximately 50, and tadpoles in batches of approximately 100. After 10 minutes, animals were transferred from the tubs back to the pens. To determine treatment effectiveness, we re-swabbed all animals or a subset (depending on the treatment) at the end of the treatment period. After the final treatment, we released all animals from the pens back into the study lakes.

#### Statistical analyses

We analyzed treatment results with linear simple and multilevel models in a Bayesian framework. All analyses except one used the brms package in R (Bürkner 2017, Bürkner 2018, R Core Team 2020). The exception was the analysis of the CMR data collected as part of the itraconazole treatments in LeConte Basin (see **Experiment-specific Methods** below). The LeConte CMR model was implemented in Stan (Carpenter et al. 2017) directly instead of via the brms interface. When using the brms package, our analysis workflow included starting with a model that included all relevant population-level (“fixed”) effects and their interactions, and checking model fit using visualizations of leave-one-out (“LOO”) probability integral transformations (Gelman et al. 2013, Vehtari et al. 2017). When suggested by the data structure or measures of model fit, we evaluated other model families or added group-level (“random”) effects to the model. We compared fits of models using LOO cross-validation and the *loo* package (Vehtari et al. 2017). For all models, we used brms defaults for priors, number of chains (4), and warmup and post-warmup iterations (1000 for each). We evaluated the adequacy of posterior samples using trace plots, Gelman-Rubin statistics (Rhat), and measures of effective sample size (“bulk-ESS”, “tail-ESS”). When using negative binomal models (most analyses), the Bd load data were rounded to integer values to produce count data.

When necessary, we developed distributional models in which predictor terms are specified for other parameters of the response distribution instead of only the mean (e.g., negative binomial overdispersion (“shape”), zero-inflation (“zi”); see brms vignette, “Estimating distributional models with brms”: https://paul-buerkner.github.io/brms/articles/brms_distreg.html). The overdispersion parameter *ϕ* controls the variance of the negative binomial distribution relative to the expected value *µ*, such that the variance of the negative binomial distribution is *µ* + *µ*^2^*/ϕ*. Modeling effects on overdispersion and zero-inflation can be important for improving model fit. For example, itraconazole treatment can reduce not only mean Bd load, but also the variation around the mean (i.e, overdispersion) and amount of zero-inflation. Improving model fit was our primary interest in using distributional models, and not gaining insights into the causes of overdispersion or zero-inflation. Therefore, when we used distributional models, we limit our descriptions of model results largely to effects of predictors on the mean.

The models described in subsequent sections are the best-fit models that resulted from the workflow outlined above. We considered predictors of group- and population-level effects and family-specific parameters to be important when the 95% credible interval (“CI”) of the estimates did not include zero, and relatively unimportant otherwise. We provide the results of all analyses in tabular form, either in the Results section for analyses describing the outcome of treatment experiments, or in **Supporting Information** for related but less central analyses. All datasets and code to replicate the analyses are available at https://github.com/SNARL1/bd-mitigation-report. To interpret the coefficients from negative binomial models, note that there is a log link for the mean (and therefore Bd load data is on a log scale). In addition, for zero-inflated negative binomial models, there is a logit link for the zero-inflation component. The key results from treatment experiments are also visualized using boxplots or dotplots. We used the former when sample sizes were relatively large and the latter when sample sizes were small and boxplots were consequently less informative. When relevant, sample sizes are displayed above the x-axis of each plot. In plots where sample sizes are displayed, the lack of sample size information for a particular group indicates that this group was intentionally not included in surveys and/or sampling. In contrast, a sample size of zero (“n=0”) indicates that this group was included in surveys and/or sampling, but that no individuals were available for capture and sampling.

### Experiment-specific methods

#### Itraconazole treatment of early life stages

Bd-caused epizootics and resulting mass die-offs of *R. sierrae* occurred in Barrett Lakes Basin during 2005 to 2007 (Vredenburg et al. 2010) and in Dusy Basin in 2009 (Jani et al. 2017). In an effort to prevent the extirpation of remnant populations, itraconazole treatments were conducted during mid-summer of 2009 in Barrett and 2010 in Dusy. Because adults typically succumb to chytridiomycosis early in an epizootic (Vredenburg et al. 2010), at the time of the experiments these populations contained primarily late-stage tadpoles, recently metamorphosed subadults, and occasionally a small number adults. We used results from basin-wide VES conducted prior to the experiments to identify the largest remaining tadpole populations, and these were selected for use in the experiments. Populations in both basins were assigned to treated and control groups at random. The Barrett experiment included three treated and three control populations, and in Dusy, where fewer frog populations remained extant, a total of three treated and two control populations were used (Table S1). For both experiments, we predicted that itraconazole treatment would reduce Bd loads and increase the survival of frogs during and after metamorphosis. In turn, this would result in more subadults counted during VES conducted in treated versus control populations in the year of and the year following treatment. Based on VES conducted before and during the treatments, for each treated population we estimate that we captured and treated 70-90% of the early life-stage animals present.

**Table 1.** Effect of itraconazole treatment in Barrett and Dusy basins on Bd loads during the following one year period (model family is zero-inflated negative binomial).

|  | Estimate | Est. Error | lo95%CI | up95%CI | Rhat | Bulk ESS | Tail Ess |
| --- | --- | --- | --- | --- | --- | --- | --- |
| **Group-level effects** |  |  |  |  |  |  |  |
| sd(Intercept) | 0.73 | 0.25 | 0.40 | 1.37 | 1.00 | 1301 | 1563 |
| **Population-level effects** |  |  |  |  |  |  |  |
| Intercept | 14.71 | 0.44 | 13.87 | 15.55 | 1.00 | 1676 | 1668 |
| overdispersion-Intercept | -1.33 | 0.08 | -1.48 | -1.18 | 1.00 | 5407 | 2754 |
| stage(tadpole) | -2.76 | 0.10 | -2.95 | -2.57 | 1.00 | 5296 | 2417 |
| basin(dusy) | -0.26 | 0.47 | -1.17 | 0.69 | 1.00 | 1966 | 2067 |
| year_std(1) | -0.40 | 0.12 | -0.63 | -0.18 | 1.00 | 3922 | 3096 |
| treatment(treated) | -1.32 | 0.50 | -2.33 | -0.32 | 1.00 | 1556 | 1614 |
| year_std(1):treatment(treated) | 1.52 | 0.18 | 1.16 | 1.87 | 1.00 | 3923 | 2645 |
| overdispersion-stage(tadpole) | 0.34 | 0.07 | 0.20 | 0.48 | 1.00 | 4540 | 3301 |
| overdispersion-basin(dusy) | 0.45 | 0.06 | 0.33 | 0.57 | 1.00 | 5399 | 3119 |
| overdispersion-year_std(1) | 0.82 | 0.07 | 0.67 | 0.96 | 1.00 | 4611 | 3192 |
| overdispersion-treatment(treated) | -0.71 | 0.07 | -0.85 | -0.57 | 1.00 | 4632 | 2889 |
| **Family-specific parameters** |  |  |  |  |  |  |  |
| zi | 0.01 | 0.00 | 0.01 | 0.02 | 1.00 | 5437 | 2653 |

In the Barrett experiment, during July 29 to August 1 we captured as many *R. sierrae* as possible (mostly tadpoles) from each pond assigned to the treated group and held them in pens. We collected skin swabs from a subset of animals to quantify Bd load. Daily itraconazole treatments were conducted during the period July 30 – August 5, and animals were treated from four to seven times, depending on the day of capture. To assess treatment effectiveness in reducing Bd loads, we collected a second set of skin swabs from a subset of animals following the final treatment. A total of 977 tadpoles and 65 subadults were treated and released back into the study ponds. In the control populations, we captured a sample of tadpoles and subadults on a single day (one population per day during August 2–4), swabbed each individual, and held them in pens until capturing was complete. Swabs were collected from a total of 75 tadpoles and 23 subadults. To quantify the longer-term effects of treatment on Bd load and frog population dynamics, we conducted post-treatment VES and swabbing at each pond in August and September 2009, and in July, August, and September 2010. In treated populations, given that we were unable to capture all animals for treatment, animals swabbed during the post-treatment period likely included a mix of treated and untreated individuals.

The Dusy experiment was identical in most respects to the Barrett experiment. During July 24–26, we captured as many animals as possible from the ponds assigned to the treated group and placed them in pens. Daily itraconazole treatments began on July 27 and lasted through August 2, resulting in seven days of treatment for all animals. In treated populations, swabs were collected from a subset of animals on July 27 before treatment began, and after the final treatment on August 2. A total of 3707 tadpoles and 125 subadults were treated. Animals in control populations were captured, swabbed, and released on July 29 (62 tadpoles and 18 subadults). We conducted follow-up VES in each pond in August and September 2010, and July and August 2011. As with the Barrett treatment, animals swabbed in treated populations during the post-treatment period likely included a mix of treated and untreated individuals.

Our analysis of the data from the experiments focused on two questions: During the one-year period following the treatments, did itraconazole treatment influence (1) Bd loads and (2) survival of treated animals? To address the first question, we developed separate models to describe (i) pre-treatment differences in Bd loads of the animals assigned to the treated and control groups, (ii) immediate effects of treatment on Bd loads, and (iii) treatment effects on post-release Bd loads in the year of treatment and the following year. Because the treatments in Barrett and Dusy Basins were virtually identical in their design, we combined the results from both experiments into a single dataset, and included basin as a predictor variable in models to account for any between-basin differences.

We evaluated pre-treatment differences in Bd load between treated and control groups using the model *bd_load ∼ (treatment x basin)* (family = negative binomial, treatment = [treated, control], basin = [Barrett, Dusy]). Life stage (tadpole, subadult) was not included in the model as a predictor because life stage and basin were collinear (i.e., most ponds were dominated by tadpoles but a few contained mostly subadults), and as such we could not estimate their separate effects. Adding a group-level effect of site_id did not improve model fit, indicating that between-pond differences were unimportant.

The immediate effect of treatment on Bd load was assessed using the model *bd_load ∼ stage + (trt_period x basin)* (family = zero-inflated negative binomial, stage = [tadpole, subadult], trt_period = [begin, end of treatment period]). We were able to include life stage in this model because many tadpoles metamorphosed into subadults during the treatment, producing a more balanced representation of life stages across sites. Plots of conditional effects suggested substantial differences in Bd load variation between life stages, treatment categories, and basins. Therefore, the overdispersion parameter was modeled as a function of all three predictor variables.

The effect of treatment on post-release Bd loads was evaluated using the model *bd_load ∼ stage + basin + (year_std x treatment) + (1 | site_id)* (family = zero-inflated negative binomial, year_std is a dummy variable in which 0 = year of treatment and 1 = year after treatment, site_id included as a group-level effect). Plots of conditional effects suggested substantial differences in Bd load variation between life stages, basins, years, and treatment groups, and therefore the overdispersion parameter was modeled as a function of all four predictor variables.

The effect of treatment on subsequent subadult counts was assessed using the model *count ∼ basin + ltadpole + (std_year x treatment) + (1 | site_id)* (family = zero-inflated negative binomial, count = number of subadults counted during a post-treatment VES, ltadpole = number of tadpoles counted (log_10_ transformed) during the same VES, site_id included as a group-level effect). The count of subadults served as a proxy for subadult survival, which could not be estimated directly (see **Methods – Microbiome augmentation of subadult frogs** for details). We included the tadpole count variable to account for differences between ponds in potential subadult production due to differences in the number of tadpoles.

During the 2010 itraconazole treatments in Dusy Basin, we tested if treatment of frogs reduced the concentration of Bd zoospores in the ponds (“zoospore pool”; Briggs et al. 2010). All associated methods and results are provided in **Supporting Information**.

#### Itraconazole treatment of adults

*LeConte Basin*. In mid-summer 2015, routine disease surveillance at one of the largest remaining Bd-naive *R. sierrae* populations detected high Bd loads and the presence of many moribund and dead frogs. In response to this epizootic, we immediately conducted two antifungal treatment experiments, one in the lower portion of the basin and one in the upper portion (Table S1). The lower basin contains two lakes and four ponds, and the upper basin contains a single lake. At the time of the experiments, all of these water bodies were occupied by *R. sierrae*. The two basins are approximately 750 m apart and are linked by streams and relatively gentle terrain; we therefore expected some movement of frogs between them.

The design of the treatment experiments in the lower and upper basins was nearly identical, differing only in the number of days spent capturing frogs for the “treated” group (three versus two days, respectively; Table S2). To simplify logistics, frogs in the treated group were captured during the first 2-3 days of the experiment, and frogs for the untreated “control” group were captured on the following day (day 3 or 4). All frogs included in the study were adults, and were collected opportunistically. Frogs that were visibly sick (as indicated by an impaired righting reflex) were excluded because these frogs were likely within hours of death. In the lower basin, a total of 359 and 102 frogs were captured for the treated and control groups, respectively. In the upper basin, these totals were 206 and 74 frogs. Although we spent 3-4 days capturing frogs for the experiments, because of the large size of this population, these totals are likely a relatively small proportion of the total frog population in the basin.

**Table 2.** Effect of itraconazole treatment in Barrett and Dusy basins on counts of subadults during the following one year period (model family is negative binomial).

|  | Estimate | Est. Error | lo95%CI | up95%CI | Rhat | Bulk ESS | Tail Ess |
| --- | --- | --- | --- | --- | --- | --- | --- |
| **Group-level effects** |  |  |  |  |  |  |  |
| sd(Intercept) | 0.33 | 0.27 | 0.01 | 1.00 | 1.00 | 1428 | 1686 |
| **Population-level effects** |  |  |  |  |  |  |  |
| Intercept | 0.76 | 0.72 | -0.57 | 2.30 | 1.00 | 2810 | 2147 |
| basin(dusy) | 0.43 | 0.50 | -0.52 | 1.43 | 1.00 | 3470 | 2893 |
| ltadpole | 0.45 | 0.24 | -0.02 | 0.91 | 1.00 | 3482 | 2531 |
| year_std(1) | -0.28 | 0.65 | -1.57 | 0.93 | 1.00 | 2734 | 2226 |
| treatment(treated) | 1.65 | 0.67 | 0.35 | 2.97 | 1.00 | 2910 | 2770 |
| year_std(1):treatment(treated) | -2.16 | 0.86 | -3.84 | -0.51 | 1.00 | 2546 | 2616 |
| **Family-specific parameters** |  |  |  |  |  |  |  |
| overdispersion | 0.65 | 0.18 | 0.37 | 1.06 | 1.00 | 4460 | 3320 |

We conducted the itraconazole treatments as described in **General Methods** and Table S2. Briefly, frogs in the treated category were swabbed, tagged with PIT tags to allow identification of individuals, measured, and weighed immediately following capture, and held in pens for the duration of the treatment period. Control frogs were captured, swabbed, PIT tagged, measured, weighed, and released. To determine treatment effectiveness, 93 frogs in the lower basin treated group (31 from each capture date) and 50 frogs in the upper basin treated group (25 from each capture date) were re-swabbed on the day prior to the final treatment. After the last treatment, all frogs were released from the pens.

To estimate pre-treatment differences in Bd loads of frogs assigned to the treated and control groups, we used the model *bd_load ∼ (location x group)* (family = negative binomial, location = [lower, upper], group = [treated, control]). To evaluate the immediate effect of treatment on Bd loads, we used the model *bd_load ∼ (location x trt_period)* (family = negative binomial, trt_period = [begin, end of treatment period]). For both analyses, we excluded any frogs that died during the treatment period. We evaluated differences in Bd loads of frogs that lived versus died during the treatment period using the model *trt_died ∼ (lbd_load x location)* (family = bernoulli, trt_died = [true, false], lbdload = log_10_(bd_load + 1) on swabs collected immediately prior to the treatment period).

The outcome of the treatment experiments was quantified using CMR surveys conducted during the summers of 2016, 2017, and 2018 (see **General Methods** for details). No post-treatment surveys were possible in 2015 because the frog active season was nearly over by the time the treatments were completed. There were 1-3 primary periods per summer, and except the final period in 2018 when only a one-day survey was conducted, all frog populations were surveyed on each of three consecutive days. Any untagged frogs captured during the surveys (i.e., frogs that were not part of the initial treatment phase of the experiment; “non-experimental”) were tagged and processed as described above.

We used open population multi-state hidden Markov models to describe subsequent population dynamics including survival and recruitment, while accounting for imperfect detection (see **Supporting Information** for details). Briefly, we estimated population size over time using parameter-expanded Bayesian data augmentation, which augments the capture histories of observed individuals with a large number of capture histories for individuals that were never detected (Royle and Dorazio 2012). The states included (1) “not recruited”, (2) “alive at the upper site”, (3) “alive at the lower site”, and (4) “dead”. On any particular survey, we considered three possible observations of an individual: (1) “alive at the upper site”, (2) “alive at the lower site”, and (3) “not detected”. The model structure builds on the work of Joseph and Knapp (2018), tracking individual Bd loads over time, allowing the expected Bd load (log_10_(Bd load + 1)) to vary as a function of treatment and time, and allowing the effect of Bd load on survival to vary as a function of treatment.

#### Treasure Lakes Basin

In July 2018, we conducted an antifungal treatment of adult *R. sierrae* during a Bd epizootic in the Treasure Lakes basin, Inyo National Forest (Table S1). Epizootics occurred in all other populations in this basin during summer 2017, and its spread to the study population in 2018 provided another opportunity to conduct an itraconazole treatment on adult frogs. However, unlike the LeConte treatment described above, this treatment was conducted as a management action instead of an experiment, primarily due to the advanced stage of the epizootic and the resulting small number of adults remaining in the population. In the VES conducted just prior to the treatment period we found only 12 adult *R. sierrae*, and captured only 28 frogs during the first day of frog collection (12 person-hours). Dividing this small population into treated and control groups would have provided little statistical power to detect between-group differences given low anticipated post-treatment recapture rates. In addition, fewer frogs would have received antifungal treatment. The lack of an experimental design limits the generality of our findings, but the treatment is nonetheless included here because of the additional insights the results provide.

We used the same methods as described for the LeConte treatments, with two important differences: (1) all frogs were treated (there was no control group), and (2) new frogs were captured from the lake and added to the pens during the first five days of the 7-day treatment period. We treated 28 frogs on July 16, then added and treated an additional 24, 7, 7, 4, and 4 frogs on July 17 through 21, respectively. Although we captured and treated a total of 74 frogs, we released only 33 live frogs at the end of the treatment due to chytridiomycosis-caused mortality throughout the treatment period. In addition to swabs collected from all frogs immediately following their initial capture, we also collected swabs from each surviving frog after the final itraconazole treatment. We compared Bd loads measured before and after treatment using the model *bd_load ∼ trt_period* (family = negative binomial, trt_period = [begin, end]). We conducted follow-up VES, swabbing, and CMR surveys one month after the 2018 treatment (August 21-23), and again in 2019 (August 15-16) and 2020 (June 23-25).

The greater range of treatment days to which frogs were exposed (compared to the LeConte treatments) provided an opportunity to evaluate the effectiveness of itraconazole treatment on Bd loads as a function of the number of daily treatments frogs received. We calculated treatment effectiveness for individual frogs as the negative log ratio of pre-treatment to post-treatment Bd loads (hereafter, “LRR”): -log_10_((load_pre_ + 1)/(load_post_ + 1). Larger absolute values of LRR indicate a larger reduction in Bd load. To evaluate the factors influencing treatment effectiveness on individual frogs, we used the model *LRR ∼ (capture_bdload_std x days_inside)* (family = gaussian, capture_bdload_std = Bd load prior to treatment standardized to mean = 0 and standard deviation = 1, days_inside = number of treatments a frog received).

#### Microbiome augmentation of subadult frogs

By 2012, the Bd epizootic in Dusy Basin (see **Methods - Itraconazole treatment of early life stages**) had caused the extirpation of most *R. sierrae* populations at this location. Extant populations contained only late-stage tadpoles and recently metamorphosed subadults, and given the absence of any adults, were presumed to represent the final cohorts at these sites. In July 2012, we initiated an experiment to test the combined effect of itraconazole treatment and *J. lividum* augmentation on Bd load and frog survival. This experiment was conducted at a single pond where late-stage tadpoles and recently-metamorphosed subadults were still relatively abundant. This pond was also used in the 2010 experiment in which early life stages were treated with itraconazole (Table S1).

In designing this experiment, we assumed that probiotic bacteria would affect Bd-frog dynamics by reducing Bd colonization of relatively lightly infected frogs, instead of by reducing Bd load on heavily infected frogs (R. Harris, personal communication). Therefore, prior to exposing frogs to *J. lividum*, we first reduced their Bd loads with a 7-day itraconazole treatment. The experiment focused solely on subadults, and included a treated group (itraconazole treatment followed by *J. lividum* exposure) and a control group (no itraconazole, no *J. lividum*). We did not test independent or interactive effects of itraconazole treatment and *J. lividum* augmentation, a decision prompted by results from our previous experiments with early life stages showing a lack of longer-term benefits of itraconazole treatment alone (see **Results**) and the limited number of subadults available at the study pond.

Subadults were captured on July 12-13 (n = 331) and assigned at random to treated and control groups at a ratio of approximately 4:1 (271 treated, 60 control). This ratio was chosen to maximize the number of subadults receiving antifungal treatment while maintaining a sufficiently large control group such that loads could be assessed with high confidence. All animals were given group-specific toe-clips as follows: (i) control = toe 2 on left front foot, and (ii) itraconazole + *J. lividum* = toe 2 on right front foot. We used toe clips because PIT tags are too large to be used with subadults. In addition, our testing of miniature numbered tags that are read visually (VI Alpha tags: Northwest Marine Technology) indicated that they were not sufficiently visible through the relatively opaque skin of subadults. Following processing, subadults in the treated and control groups were held in separate mesh pens. To determine Bd loads prior to the start of the itraconazole treatment period, a subset of subadults from both groups were swabbed on July 12, and all control animals were released back into the study pond on July 13.

Itraconazole treatments were conducted daily on July 12-18. The number of animals in each group declined somewhat during this period due to Bd-caused mortality and occasional predation by gartersnakes. To assess the effectiveness of itraconazole treatment in reducing Bd loads over the 7-day treatment period and to quantify the amount of *J. lividum* present naturally on subadults in this population, we swabbed a subset of animals on July 19 immediately prior to *J. lividum* exposure. To compare pre-treatment Bd loads on frogs assigned to the treated and control groups, we used the model *bd_load ∼ expt_trt* (family = negative binomial, expt_trt = [treated, control]). The effectiveness of the itraconazole treatment was assessed with the model *bd_load ∼ days* (family = zero-inflated negative binomial, days = -7 (before treatment) and 0 (after treatment)). *J. lividum* for use in the experiment was obtained from the skin of an adult *R. sierrae* in Dusy Basin in 2009 and cultured using standard methods (Harris et al. 2009). On July 19, a concentrated solution of *J. lividum* culture was transported into Dusy Basin on foot in an insulated container (the insulated container ensured that the solution would remain cold during transport). On July 19-20, we bathed the itraconazole-treated subadults (n = 256) in a solution of *J. lividum* culture for 4-4.5 hours (75 and 150 mL of *J. lividum* culture per liter of lake water on July 19 and 20, respectively; concentration of *J. lividum* is unknown). At the conclusion of the second *J. lividum* bath, all animals were released back into the pond. The solution of *J. lividum* culture was carried out of the backcountry and disposed of.

To assess the longer-term effects of the combined itraconazole-*J. lividum* treatment, we surveyed the study population during the summers of 2012 (n = 3 surveys), 2013 (n = 3), 2014 (n = 1), and 2019 (n = 1). During each of these surveys, we conducted VES and captured and swabbed as many subadult frogs as possible (no adults were captured), and recorded the toe-clip (if present) for each individual. The concentration of *J. lividum* on frogs was assessed from skin swabs using qPCR (see **Supporting Information** for details).

We analyzed the collected data to determine whether subadults exposed to the combined itraconazole-*J. lividum* treatment had (1) higher concentrations of *J. lividum* and lower Bd loads than untreated control animals and non-experimental (“wild”) animals, and (2) higher survival than control animals. All analyses focused on data collected during the two months immediately following the 2012 treatment. Recaptures of control animals quickly declined to near zero, thereby precluding formal comparisons of *J. lividum* and Bd load in the treated versus control groups. We were able to compare treated versus wild frogs, but importantly, unlike the treated and control groups that each contained a single cohort of toe-clipped animals that was repeatedly sampled over time, membership of animals in the wild group changed over time as new individuals entered the group following metamorphosis and previously-metamorphosed individuals died. Given this limitation, we describe the *J. lividum* concentration and Bd load on treated versus control animals graphically only. We analyzed the *J. lividum* concentration on treated versus wild frogs using the model *jliv ∼ (days x frog_group)* (family = negative binomial, days = days since *J. lividum* exposure, frog_group = [treated, wild]). To describe Bd loads of treated versus wild frogs, we used the model *bd_load ∼ (days x frog_group)* (family = zero-inflated negative binomial).

For the second question, we used the percent of animals in each group that were recaptured as a proxy for survival. Formal assessment of the effect of treatment on survival was again not possible due to the rapid disappearance of animals in the control group, so the results are described graphically only.

## Results

### Itraconazole treatment of early life stages

In the Barrett and Dusy experiments, immediately before itraconazole treatments began, Bd loads of animals in ponds assigned to the treated and control groups were similar (Figure 1: Week -3 and -1). Model results (Table S3) confirmed that Bd load did not differ between treatment groups. In addition, basin had a weak effect (loads were lower in Dusy than Barrett), and the (treatment x basin) interaction term was unimportant, indicating that the patterns of Bd load between treated and control groups were similar in both basins.

**Fig. 1.**
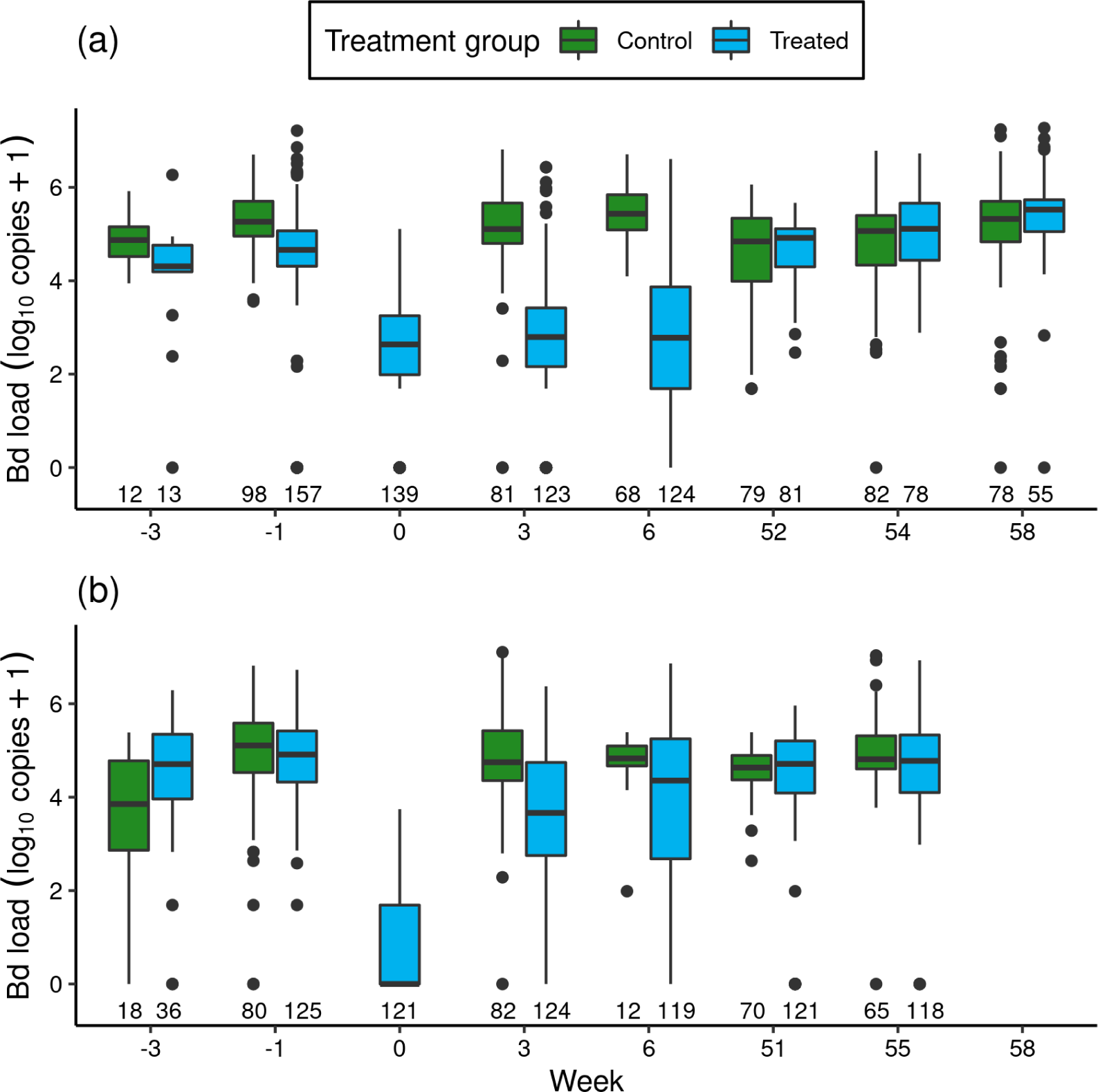
For the itraconazole treatment experiment in Barrett (a) and Dusy (b) basins, temporal patterns of Bd loads of early life stage *R. sierrae* in populations assigned to control and treated groups. Weeks -3 and -1 are pre-treatment, week 0 is the end of treatment, and weeks 3-58 are post-treatment. In the boxplots, the horizontal bar is the median, hinges represent first and third quartiles, whiskers extend to the largest and smallest values within 1.5x interquartile range beyond hinges, and dots indicate values outside the 1.5x interquartile range. The number of swabs collected in each week is displayed above the x-axis.

**Table 3.** Effect of number of days since *J. lividum* exposure and frog group (treated, wild) on *J. lividum* concentration on frogs (model family is negative binomial).

|  | Estimate | Est. Error | lo95%CI | up95%CI | Rhat | Bulk ESS | Tail Ess |
| --- | --- | --- | --- | --- | --- | --- | --- |
| **Population-level effects** |  |  |  |  |  |  |  |
| Intercept | 7.27 | 0.33 | 6.68 | 7.95 | 1.00 | 2944 | 2565 |
| days | -0.14 | 0.01 | -0.16 | -0.12 | 1.00 | 2818 | 2525 |
| frog_group(wild) | -0.29 | 0.75 | -1.71 | 1.24 | 1.00 | 2222 | 2413 |
| days:frog_group(wild) | 0.02 | 0.02 | -0.03 | 0.06 | 1.00 | 2187 | 2082 |
| **Family-specific parameters** |  |  |  |  |  |  |  |
| overdispersion | 0.33 | 0.04 | 0.26 | 0.41 | 1.00 | 3492 | 2373 |

The treatments reduced Bd loads by 1.8 orders of magnitude in Barrett and 6.1 orders of magnitude in Dusy (Figure 1: Week -1 versus 0). Model results (Table S4) substantiated the important effect of treatment period (“trt_period”; lower after treatment than before treatment). In addition, important effects on Bd load were also evident for frog life stage (lower in tadpoles than subadults), basin (higher in Dusy than Barrett), and the (basin x trt_period) interaction term. The importance of the interaction term indicated that loads were higher in Dusy than Barrett at the beginning of the treatment, but lower in Dusy than Barrett at the end of treatment (Figure 1). Finally, life stage, treatment period, and basin all had important effects on the overdispersion parameter (Table S4; Bd load was more variable in subadults than tadpoles, at the beginning than the end of the treatment period, and in Dusy than Barrett).

**Table 4.** Effect of number of days since *J. lividum* exposure and frog group (treated, wild) on Bd load on frogs (model family is zero-inflated negative binomial).

|  | Estimate | Est. Error | lo95%CI | up95%CI | Rhat | Bulk ESS | Tail Ess |
| --- | --- | --- | --- | --- | --- | --- | --- |
| **Population-level effects** |  |  |  |  |  |  |  |
| Intercept | 8.42 | 0.23 | 7.97 | 8.89 | 1.00 | 3608 | 3126 |
| days | 0.08 | 0.01 | 0.07 | 0.09 | 1.00 | 3854 | 2537 |
| frog_group(wild) | 6.74 | 0.47 | 5.86 | 7.68 | 1.00 | 2559 | 2131 |
| days:frog_group(wild) | -0.09 | 0.01 | -0.11 | -0.07 | 1.00 | 2567 | 2040 |
| **Family-specific parameters** |  |  |  |  |  |  |  |
| overdispersion | 0.56 | 0.05 | 0.47 | 0.66 | 1.00 | 3773 | 2677 |
| zi | 0.08 | 0.02 | 0.05 | 0.12 | 1.00 | 4066 | 2525 |

After release of the treated animals back into the study ponds, the reduction in Bd load in treated versus control groups that was evident at the end of the treatment period persisted for at least the next 1.5 months (Figure 1: Week > 0). Results from a model of predictors of Bd load over the 1-year post-release period (Table 1) showed important effects on Bd load of most predictor variables, including treatment (treated lower than control), life stage (lower in tadpoles than subadults), year (lower in the year following treatment (year 1) than the year of treatment (year 0)), and the (year x treatment) interaction term. Basin did not have an important effect. The (year x treatment) term indicated that Bd loads were lower in the treated group than the control group in year 0, but by year 1 loads in the treated group had increased such that Bd loads of the treated and control groups were similar. Therefore, although the treatment effect was evident for more than a month, Bd loads on animals in treated populations returned to pre-treatment levels in the year following treatment (Figure 1). All predictors of the overdispersion parameter had important effects.

The reduction in Bd load caused by the treatment was associated with increased counts of subadults in treated versus control populations (Figure 2). Model results (Table 2) indicated that treatment and the (year x treatment) interaction term had important effects. The effects of tadpole count, basin, and year were unimportant. The interaction term indicated that treated populations had higher subadult counts than control populations in the year of the treatment, but that counts in treated populations in the year following treatment were low and similar to those in control populations (Figure 2). Therefore, mirroring the longer-term effects of treatment on Bd load, the increase in subadult counts in treated populations in the 1.5 months following treatment was no longer evident in the year following treatment.

**Fig. 2.**
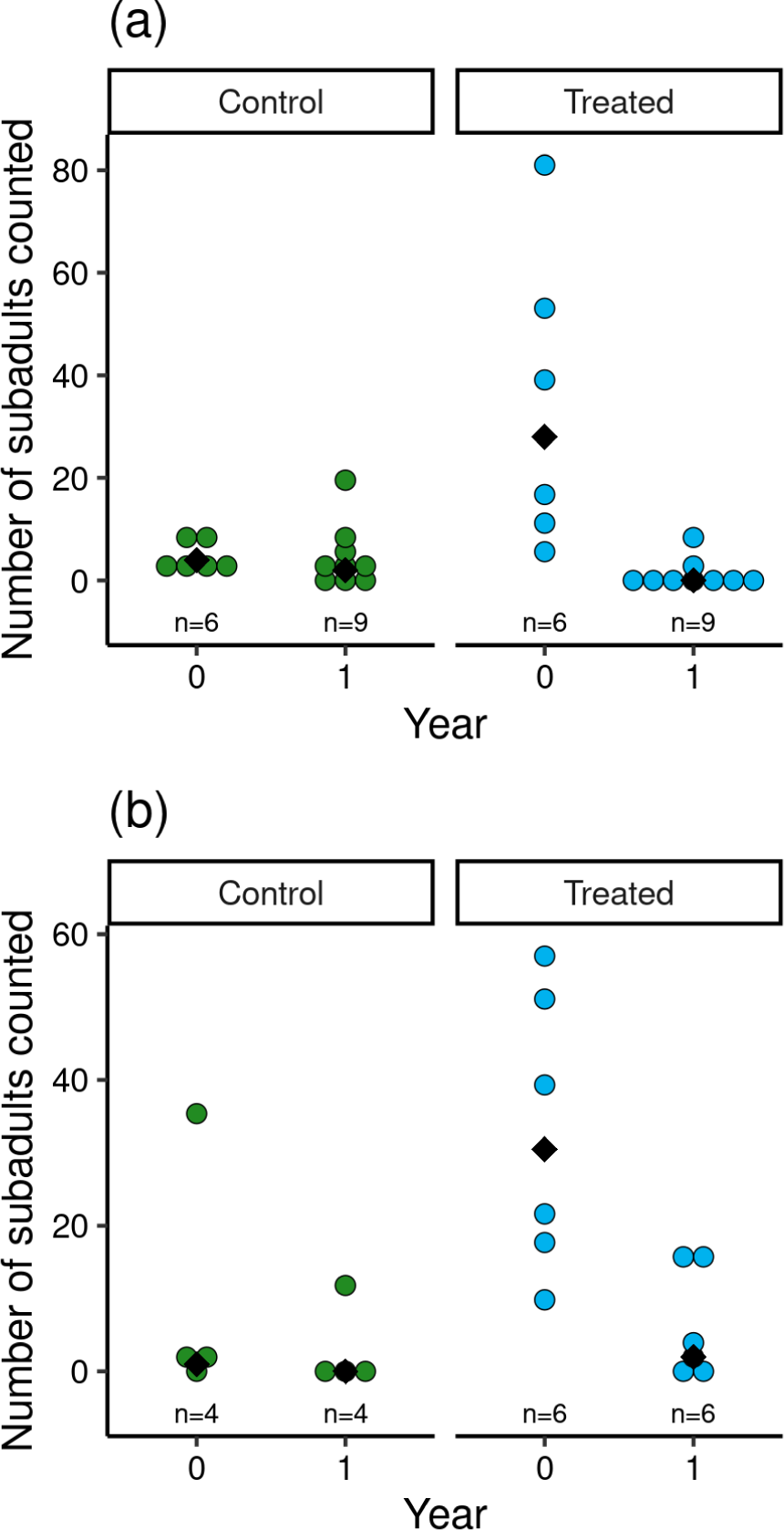
For control and treated populations in Barrett (a) and Dusy (b) basins, post-treatment counts of *R. sierrae* subadults in the year the treatment was conducted (year = 0) and the year following the treatment (year = 1). Each dot indicates the count made during a survey of one of the study ponds, and median values for each treatment group are indicated with a black diamond. The total number of surveys is displayed above the x-axis.

### Itraconazole treatment of adults

#### LeConte Basin

In the two itraconazole treatment experiments conducted in LeConte Basin, prior to the treatment period, adult *R. sierrae* assigned to the treated and control groups had very high Bd loads, above the level at which symptoms of severe chytridiomycosis are evident (Figure 3). Bd loads in the control group were somewhat higher than in the treated group, likely because control frogs were captured and processed 1-3 days later than frogs assigned to the treated group (Table S2) and during a period when Bd loads were increasing in the study populations. Model results (Table S6) affirmed an important pre-treatment difference in Bd load between treatment groups (treated groups lower than control groups). Location and the (treatment x location) interaction term were both unimportant, with the latter indicating that the pattern of Bd load between treatment groups was similar in the lower and upper basins.

**Fig. 3.**
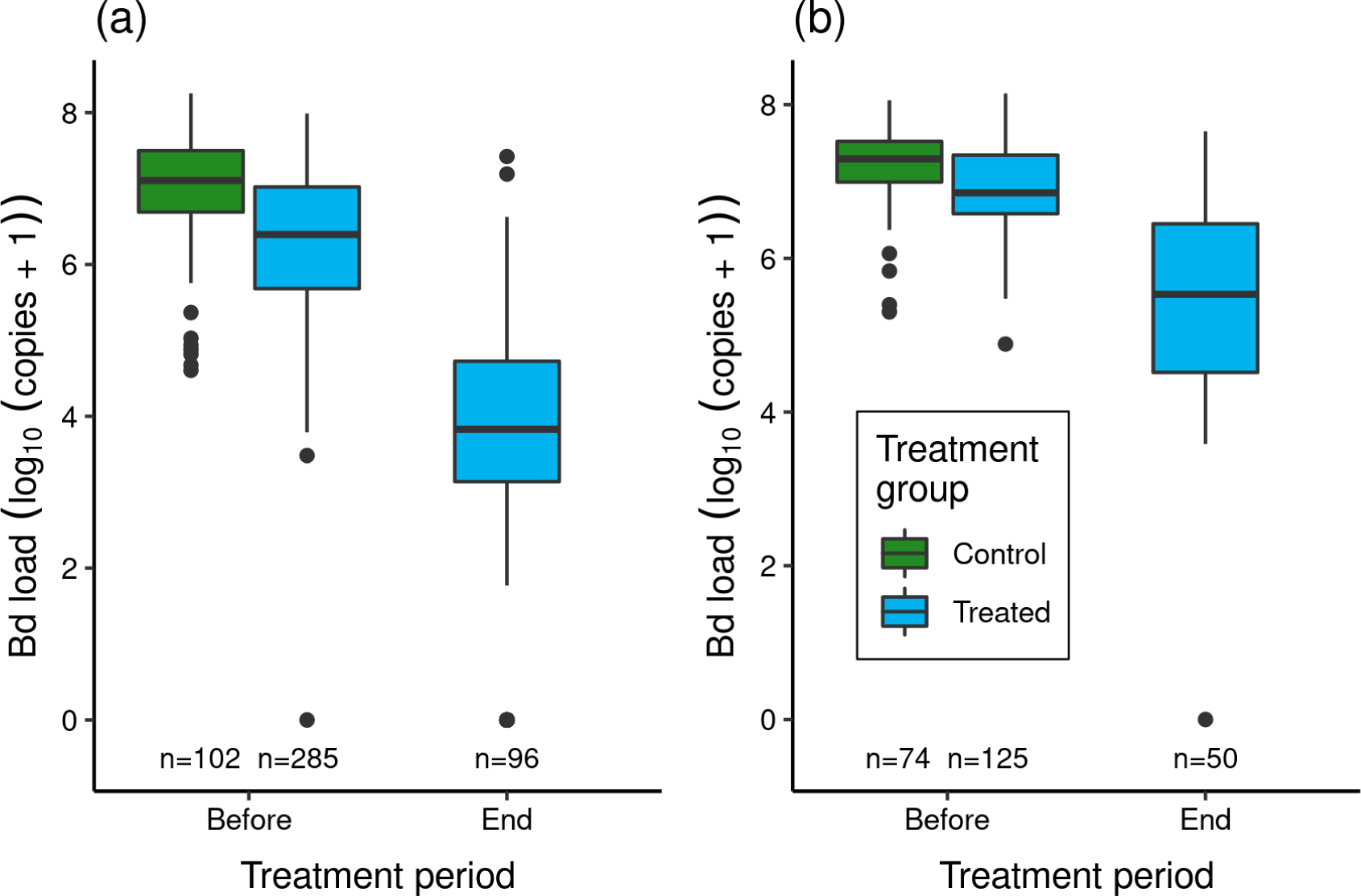
Effect of itraconazole treatment on Bd loads of adult *R. sierrae* in the 2015 LeConte treatment experiment: (a) lower basin, and (b) upper basin. The legend for both panels is provided in (b). Box plots show Bd loads on frogs in the control (untreated) and treated groups before the treatment began and at the end of the treatment period. Control frogs were processed and released before the treatment period, and therefore no Bd samples were collected from control frogs at the end of this period. Only frogs that survived to the end of the treatment period and were released back into the study lakes are included. The number of swabs collected from frogs in each category are displayed above the x-axis. Box plot components are as in Figure 1.

Samples collected one day prior to the end of the 8-9 day treatment period (Table S2) indicated that in both experiments the treatment reduced Bd loads on treated frogs by 1.4-2.7 orders of magnitude (Figure 3). Model results (Table S7) corroborated the important effect of treatment on Bd load. The effect of location was also important (higher in the upper than lower basin), as was the (location x trt_period) interaction term, with the latter indicating that Bd loads before and at the end of treatment were both higher in the upper than lower basin.

During the treatment period, 74 of the lower basin treated frogs and 80 of the upper basin treated frogs died. All control frogs survived during the several hour period between capture, processing, and release. Of the treated frogs that died, most did so during the first half of the treatment period (lower basin: 73%; upper basin: 74%), consistent with frogs succumbing to chytridiomycosis. However, Bd load was not an important predictor of whether frogs died versus survived (Table S8). Location and the (location x bd_load) interaction term were also unimportant.

Based on CMR modeling, across the entire duration of the experiment (2015-2018), the 1206 unique individuals included in the study were estimated to represent approximately 80% (posterior median) of the adults that existed in the LeConte population during this time (CI: 75% – 88%). Between frog release in 2015 and the final survey in 2018, seven recaptured individuals moved between the two basins. All seven were in the treated group and moved from the upper to the lower basin. These individuals were included in counts of unique individuals in the basin in which they were captured. In CMR surveys conducted during 2016-2018, a total of 2208 adult frogs were captured, representing 831 unique individuals. Of the 745 unique frogs captured in the lower basin, 132 were in the treated group, two were in the control group, and 611 were not part of the original treatment experiment (“non-experimental” frogs). In the upper basin, 89 unique frogs were captured, of which 81 were in the treated group and eight were non-experimental. No control frogs were captured in the upper basin. In total, during the three year post-treatment period across both experiments, 54% of treated frogs and 1% of control frogs were recaptured. The 619 non-experimental frogs could have either survived the 2015 Bd epizootic as adults or recruited into the adult population after the epizootic. Frogs in this group spanned a broad range of sizes (40–75 mm, median = 50 mm), and 83% were larger than the 40–45 mm range that characterizes recent adult *R. sierrae* recruits. In addition, all of the larger untagged frogs (> 45 mm) were captured in 2016, and all frogs captured in 2017 and 2018 were in the 40–45 mm range.

Importantly, the reduced loads of the treated group after the 2015 treatment period were maintained in all three post-treatment years (Figure 4a). Bd load dynamics in control frogs are less clear because only two control frogs were recaptured during 2016-2018. However, these two recaptured control frogs were recaptured in the year after the treatments (2016), and both had relatively low loads in 2015 relative to the rest of the individuals in the control group (Figure 4a). During the 2016-2018 period, Bd loads in the non-experimental group were relatively low and similar to those of the treated group.

**Fig. 4.**
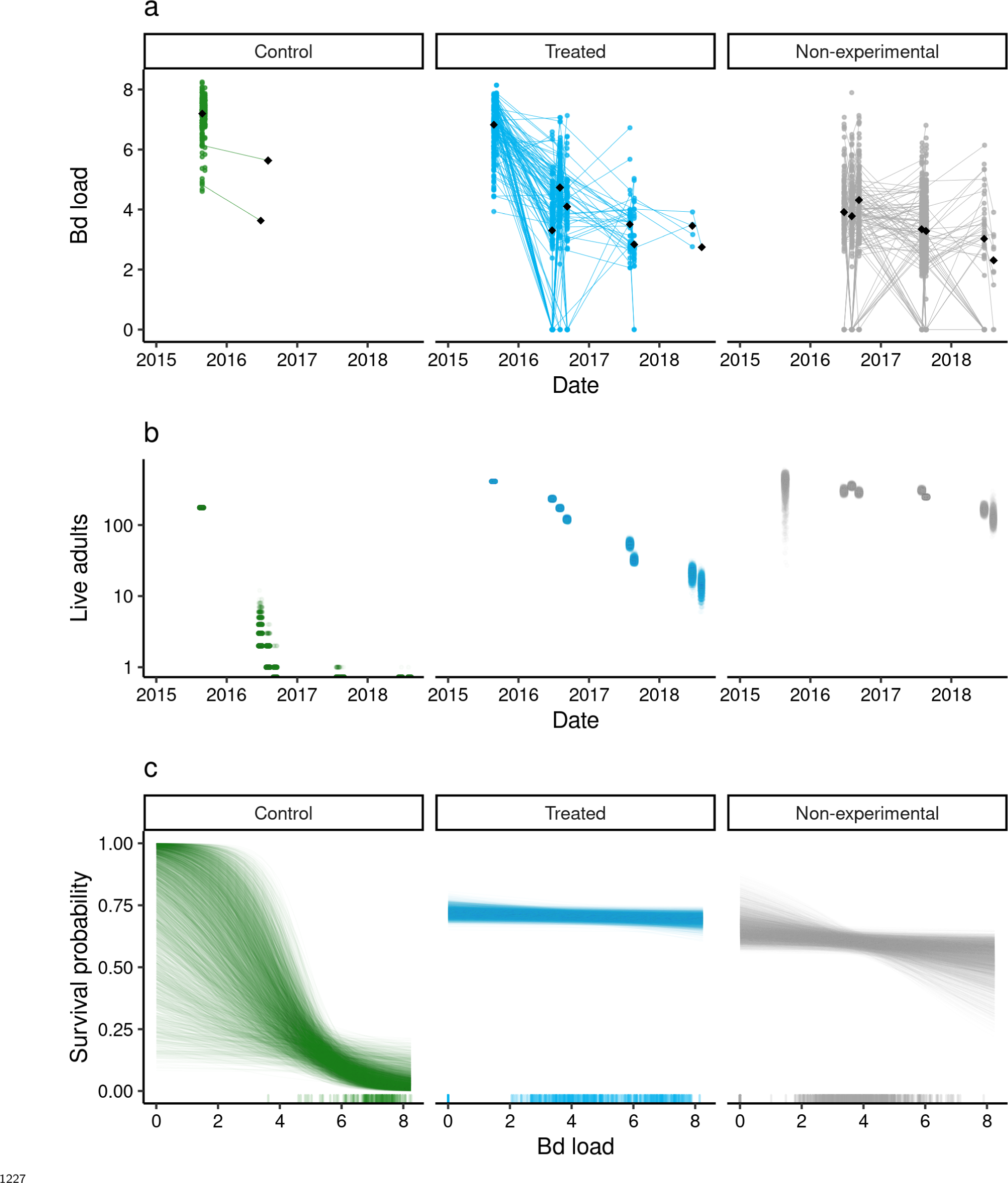
Outcome of the LeConte treatment experiment with adult *R. sierrae*, showing results for control, treated, and non-experimental animals. Time series from 2015 to 2018 of observed (a) Bd loads, with lines connecting sequential observations of tagged individuals, (b) posterior estimates for the number of live adults (abundance) in each group, where each point is a draw from the posterior, and (c) estimated relationships between Bd load and adult survival probability during the entire study period, with one line for each posterior draw. A rug along the x-axis displays the observed distributions of Bd load. In (a) and (b), the date tick marks indicate January-01 of each year. In (a) and (c), the Bd load axis shows Bd loads as log_10_(copies + 1).

Overall, the LeConte adult population declined in abundance from 2015 to 2018, with the most rapid declines in the control group (Figure 4b). Between the end of the treatments in 2015 and the first surveys of 2016, the number of animals surviving in the control group dropped from 176 to 8 (CI: 2 – 18). By the end of the summer in 2016, the posterior median for the number of surviving control animals was 0 (CI: 0 – 0). The rate of decline was slower in both the treated and non-experimental groups (Figure 4b). Nonetheless, despite the treatment, by 2018 the study populations in the lower and upper basins had declined to few remaining adults. In the last primary period of 2018, the posterior median for the number of treated frogs alive across both basins was 9 (CI: 3 – 17), 125 for non-experimental frogs (CI: 87 – 188), and zero for controls (CI: 0 – 0).

Interestingly, Bd load had a stronger negative effect on survival in the control group relative to the treated and non-experimental groups (posterior probabilities = 0.99 and 0.99, for control versus treated, and control versus non-experimental, respectively; Figure 4c). This was the case despite considerable overlap in Bd loads between the control and treated/nonexperimental groups (Figure 4a, 4c).

Detection probabilities in the study populations varied over time, but overall were comparable to estimates previously reported from other populations (Joseph and Knapp 2018). The primary period with the highest detection probabilities had a posterior median detection probability of 0.52 (CI: 0.49, 0.56). In contrast, the primary period with the lowest detection probabilities had a posterior median of 0.1 (CI: 0.05, 0.17). On an average primary period, posterior median detection probability was 0.28 (CI: 0.17, 0.44).

#### Treasure Lakes Basin

Similar to the situation in LeConte Basin, adult frog Bd loads were very high at Treasure Lake during early summer 2018, and at the start of the itraconazole treatment (Figure 5). Itraconazole treatment reduced Bd loads by more than two orders of magnitude (Figure 5; Bd loads on 2018-07-23 versus on days 2018-07-16 to 2018-07-21). Model results affirmed the important effect of treatment (Table S9).

**Fig. 5.**
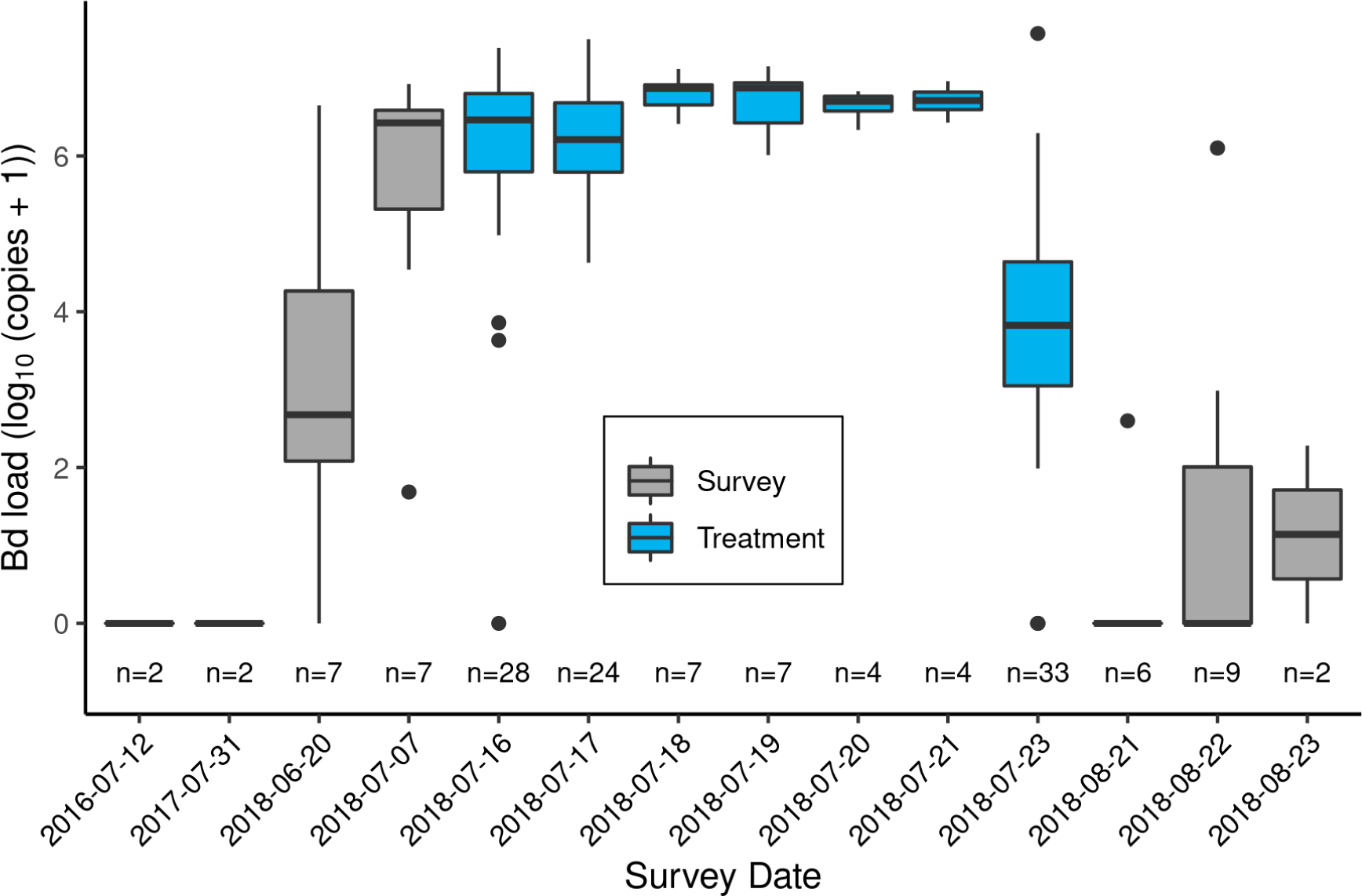
Bd loads for adult *R. sierrae* at the Treasure Lake study site before the Bd epizootic (2016-2017), and throughout the 2018 summer when the Bd epizootic began and the antifungal treatment occurred. Box colors indicate Bd loads from pre- and post-treatment periods (gray) and during the treatment (blue). The number of swabs collected on each date is displayed above the x-axis. Box plot components are as in Figure 1. During the treatment period, individual frogs were swabbed for Bd immediately following their initial capture and again just prior to their release (on 2018-07-23).

The number of itraconazole treatments a frog received (“days_inside”) increased treatment effectiveness (Table S10). Initial Bd load (“capture_bdload_std”) and the (days_inside x capture_bdload_std) interaction term were both unimportant (Table S10). Of the 33 frogs that were released back into the lake following treatment, 16 were recaptured in the CMR survey conducted one month later (Figure 5). In addition, one non-experimental adult frog was captured, and one dead tagged (i.e., treated) adult was found. Bd loads of most recaptured frogs were low compared to those of frogs at the start and end of the treatment period (Figure 5, S3). There was no obvious relationship between the number of treatments a frog received and whether or not it was recaptured one month later (Figure S3). In surveys conducted in 2019 (the year following treatment) and 2020, we observed no *R. sierrae* of any life stage. Therefore, despite the substantial reduction in Bd loads caused by the 2018 treatment and the relatively large fraction of treated frogs recaptured one month later, few or no frogs survived overwinter until summer 2019.

### Microbiome augmentation of subadult frogs

In the 2012 Dusy Basin microbiome augmentation experiment, prior to the itraconazole treatment, Bd loads were similar in subadults assigned to the control and treated groups (Figure 6: day = -7). Model results (Table S11) affirmed that pre-treatment Bd loads of the two groups were not different. Itraconazole treatment reduced Bd loads almost four orders of magnitude (Figure 6: day -7 versus 0), and model results (Table S12) substantiated this important effect.

**Fig. 6.**
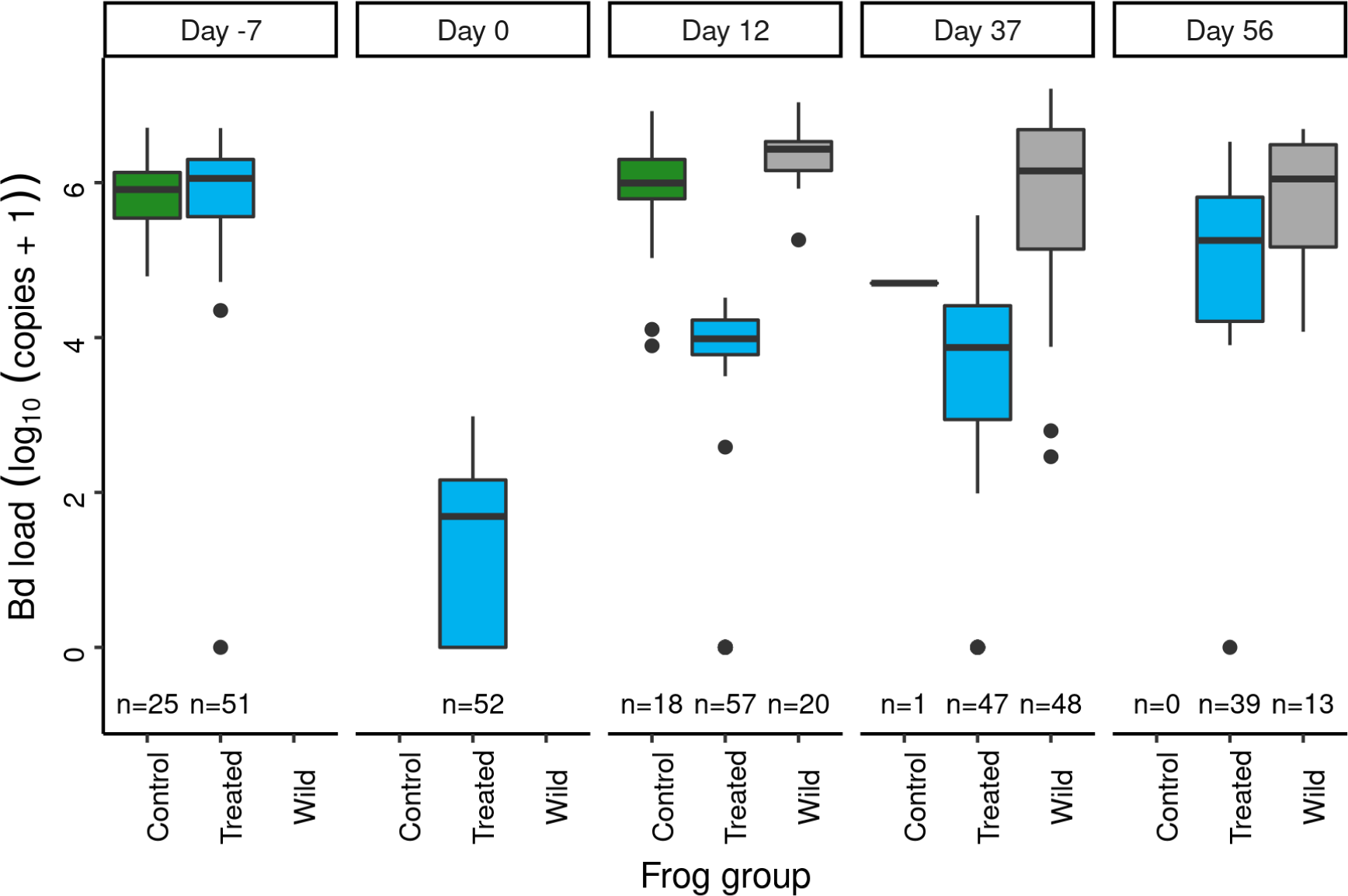
In the Dusy Basin *J. lividum* augmentation experiment, temporal patterns of Bd loads on subadult *R. sierrae* in the control, treated, and wild groups. Panel labels indicate the number of days since *J. lividum* exposure. Prior to the exposure of frogs in the treated group to *J. lividum* on days 0 and 1, frogs in the treated group were treated with itraconazole on days -6 to -1 to reduce their Bd loads. The number of swabs collected on each day is displayed above the x-axis.

*J. lividum* exposure started on the day following the last day of itraconazole treatment, and on that day *J. lividum* concentrations on subadults assigned to the treated group were either zero or near-zero for all individuals (day 0; Figure 7). Twelve days after the two *J. lividum* baths to which animals in the treated group were subjected, *J. lividum* concentrations on subadults were high, but unexpectedly the concentrations were similarly high in treated, control, and wild groups instead of only in the treated group (day 0 versus 12; Figure 7). This is consistent with transfer of *J. lividum* from treated animals to other frogs in the pond that had not been treated with itraconazole or bathed experimentally in high concentrations of *J. lividum*. However, over the following two months, *J. lividum* concentrations on subadults in all three groups declined to near baseline levels (Figure 7). Formal comparison of *J. lividum* concentrations from day 12 to day 56 across all three groups was not possible due to the almost complete absence of control frogs on days 37 and 56. However, a model that included the treated and wild groups indicated an important negative effect of the number of days since *J. lividum* exposure on *J. lividum* concentration, but no effect of group or the (day x group) interaction term (Table 3). Therefore, *J. lividum* concentrations declined over the 2-month period and at similar rates in both treated and wild frogs.

**Fig. 7.**
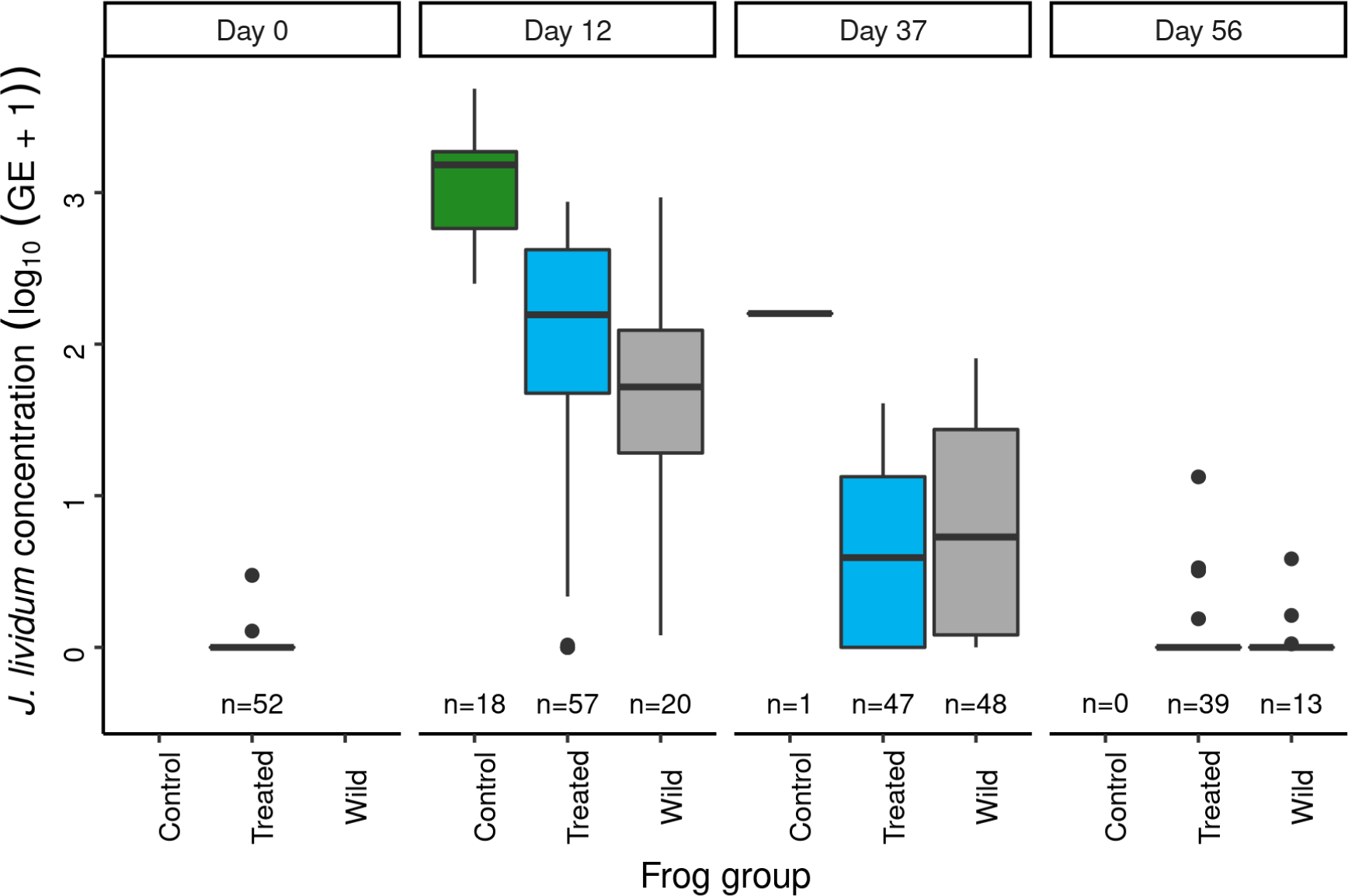
In the Dusy Basin *J. lividum* augmentation experiment, temporal patterns of *J. lividum* concentrations on subadult *R. sierrae* in the treated, control, and wild groups. Panel labels indicate the number of days since *J. lividum* exposure. Prior to the exposure of frogs in the treated group to *J. lividum* on days 0 and 1, frogs in the treated group were treated with itraconazole on days -6 to -1 to reduce their Bd loads. *J. lividum* concentrations on day 0 are from samples collected from frogs in the treated group just prior to the first *J. lividum* exposure. The number of swabs collected on each day is displayed above the x-axis.

Following release of frogs in the treated group (itraconazole-treated and *J. lividum*-exposed) back into the study pond, their Bd loads increased steadily and reached pre-treatment levels after two months (Figure 6: day 56). Due to the rapid loss of frogs in the control group, formal comparison of Bd loads from day 12 to day 56 across all three groups was not possible. However, a model that included the treated and wild groups indicated that Bd loads increased during this period, and that Bd loads of wild frogs were higher than those of treated frogs (Table 4). In addition, there was an important effect of the (days x group) interaction term, due to increasing loads of treated frogs versus relatively constant loads of wild frogs. Together, these results indicate that the combined effect of itraconazole treatment and *J. lividum* exposure was ineffective in preventing the increase in Bd loads to pre-treatment levels. Increased concentrations of *J. lividum* on control and wild frogs also had no obvious effect on Bd load.

During the three surveys of the study population conducted in 2012, the percent of frogs in the treated and control groups that were recaptured declined, but the rate of decline was steeper in the control versus treated group (Figure 8). Although no formal analysis is possible due to the relatively few sample points, the results suggest that the itraconazole-*J. lividum* treatment increased frog survival over the two month period following treatment. Nonetheless, during surveys conducted in 2013 (one year after treatment), only a single experimental (i.e., toe-clipped) animal was captured. This animal was detected during a survey conducted in early summer, and was a member of the treated group. No *R. sierrae* of any life stage were detected during surveys in 2014 and 2019. In conclusion, the combined itraconazole treatment and *J. lividum* exposure did not protect frogs against Bd infection and increase survival sufficiently to allow persistence of this population over the longer-term.

**Fig. 8.**
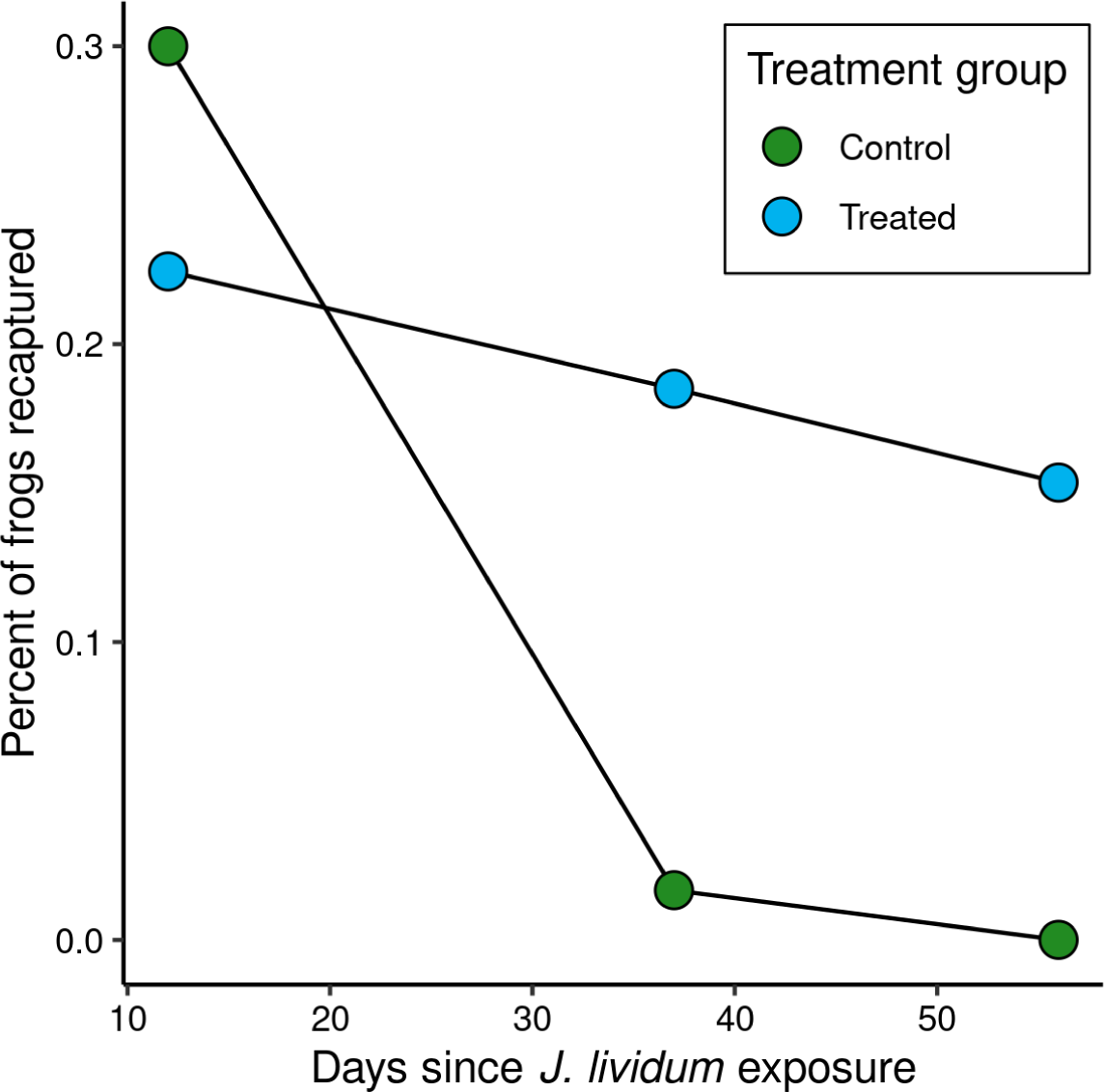
In the Dusy Basin *J. lividum* augmentation experiment, the percent of frogs in the treated and control groups recaptured during the two months following *J. lividum* exposure. The number of subadults captured on each survey is given in Figure 8 (number of subadults = number of swabs).

## Discussion

The devastating effect of chytridiomycosis on amphibian populations worldwide (Scheele et al. 2019) highlights the need for effective strategies to mitigate disease impacts in the wild following Bd emergence. Potential strategies include those aimed at eliminating Bd or facilitating host-pathogen coexistence (Garner et al. 2016). Complete eradication of Bd from amphibians and the environment will rarely be feasible, but in an isolated and simple ecosystem, treatment of all amphibians and chemical disinfection of aquatic habitats eliminated Bd over the long term (Bosch et al. 2015). Similar methods applied to a more complex system failed to achieve long-term eradication of Bd (Fernández-Loras et al. 2020).

Although many examples exist of Bd infection and resulting disease driving amphibian hosts to extinction or near-extinction (Scheele et al. 2019), amphibian-Bd coexistence is also possible. Coexistence can result from multiple mechanisms (Brannelly et al. 2021). Of these, wildlife disease management interventions often target density-dependent transmission and host resistance/tolerance (Woodhams et al. 2011, Garner et al. 2016). Reducing amphibian density or treating amphibians with antifungal agents could reduce density-dependent transmission rates, and result in increased survival and population persistence in an enzootic state (Briggs et al. 2010). In immunocompetent species and life stages (Rollins-Smith 1998, Grogan et al. 2018b), treatment could increase host resistance or tolerance by reducing Bd growth and allowing the full development of adaptive immunity, possibly increasing survival and population persistence (Woodhams et al. 2011). Host resistance or tolerance might also be increased by augmenting the microbiome on amphibian skin with antifungal probiotics that provide at least partial protection against Bd infection (Harris et al. 2009, Bletz et al. 2013).

Numerous field trials have been conducted to test these possibilities, but most have failed to increase the long-term persistence and growth of affected amphibian populations (Garner et al. 2016). Treatment of larval or post-metamorphic frogs with antifungal drugs has been attempted in several species, and generally produces short-term reductions in Bd prevalence or load and, in some cases, increases in frog survival (Hardy et al. 2015, Hudson et al. 2016, Geiger et al. 2017). However, these effects typically disappear within weeks or months of treatment, with little consequence for long-term population persistence (but see Hardy et al. 2015 for a possible exception). Laboratory experiments in which the frog skin microbiome was augmented with antifungal probiotics have demonstrated the potential of this method (Harris et al. 2009, Kueneman et al. 2016), but field trials are lacking.

The six field trials we conducted were ineffective in facilitating long-term frog-Bd coexistence. This difficulty in altering frog-Bd dynamics is consistent with theoretical work that suggests a narrow range of parameter space within which treatment strategies might prevent Bd-driven host extinctions (Drawert et al. 2017). Nonetheless, compared to disinfecting the environment or reducing host density, the treatment of infected amphibians with antifungal agents is predicted to have the greatest likelihood of a beneficial outcome and the lowest risk of reducing population persistence (Drawert et al. 2017), and is therefore worthy of detailed evaluation. In addition, carefully-designed field experiments can provide important insights into host-Bd dynamics relevant to disease mitigation efforts that generally cannot be gained from observational or theoretical studies alone. In the following sections we summarize the key results from our treatments and highlight the likely causes of the failure to facilitate long-term population persistence.

### Itraconazole treatment of early life stages

In the two itraconazole treatment experiments that focused on early life stage *R. sierrae* (Dusy and Barrett basins), Bd loads were reduced in treated populations and remained relatively low during the summer in which the treatment was conducted. In addition, this reduction in Bd load was associated with increased survival of subadults (as measured by increased numbers of subadults counted during VES). However, treatment effects were short-lived. Within a year, Bd loads returned to the high levels characteristic of the control populations, and subadult survival was reduced to zero or near-zero. In addition, the short-term increase in subadult survival did not increase recruitment into the adult population, and all control and treated populations in both basins were eventually extirpated. The disappearance of any treatment effect within one year is similar to that reported by Geiger et al. (2017) following antifungal treatment of tadpoles of the common midwife toad (*Alytes obstetricans*), and is consistent with tadpoles and subadults having relatively low immunocompetence (Bakar et al. 2016, Grogan et al. 2018b). Hardy et al. (2015) treated recently-metamorphosed Cascades frogs (*Rana cascadae*) with itraconazole, and reported increased survival of treated animals the following year. This relatively long-term benefit of antifungal treatment was not observed in either of our early life stage treatments, perhaps indicative of variation in immunocompetence of this life stage between even closely-related species.

### Itraconazole treatment of adults

In contrast to the relatively short-lived effects when early life stages were treated, the two itraconazole treatment experiments in LeConte that focused on *R. sierrae* adults reduced Bd load and increased frog survival throughout the three-year post-treatment period of the study. The reduced Bd loads on treated adults were similar in magnitude to those of adults in persistent MYL frog populations characterized by enzootic Bd dynamics (Briggs et al. 2010, Knapp et al. 2011, Joseph and Knapp 2018). In addition to increased resistance, the results also indicate that treated frogs had higher tolerance of Bd infection (see Schneider and Ayres (2008) and Soares et al. (2017) for recent reviews of resistance, tolerance, and the role of adaptive immunity in both). Specifically, although treated frogs had lower loads than control frogs throughout the post-treatment period, Bd load distributions for frogs in the two groups nonetheless overlapped. In the region of overlap (3.6–7.6 copies), survival probability was near zero for control frogs, but was higher and relatively constant for treated frogs. This higher survival at a given Bd load value is consistent with increased tolerance of Bd infection and, along with increased resistance, is suggestive of treated frogs having mounted an effective adaptive immune response against Bd (McMahon et al. 2014, Ellison et al. 2015, Grogan et al. 2018a).

Despite the extended period of reduced Bd loads and increased frog survival, the adult population declined in each of the post-treatment years, and by 2018 few treated frogs remained. This decline was likely due to a combination of reduced adult survival and insufficient recruitment of new adults. Survival of treated adults, although higher than that for control frogs (annual survival for 2015-2016, 2016-2017, and 2017-2018 = 0.56, 0.17, and 0.31, respectively), was generally still lower than that of most persisting enzootic MYL frog populations (Briggs et al. 2010, Joseph and Knapp 2018). Regarding recruitment, we counted hundreds of tadpoles and subadults during VES conducted during each of the post-treatment years (maximum counts during 2016, 2017, and 2018 = 614, 2434, and 480, respectively), but captured relatively few new adult recruits during the same period. Of the 102 untagged (i.e., non-experimental) adults we captured that had sizes typical of new recruits (40–45 mm), 91 were tagged in 2016, nine in 2017, and two in 2018. The low recruitment of new adults in 2017 and 2018 despite large numbers of early life stages resembles recruitment levels we have observed in other enzootic *R. sierrae* populations, and is likely a consequence of high chytridiomycosis-caused mortality of frogs during and soon after metamorphosis (Joseph and Knapp 2018, and results from Barrett and Dusy treatments described above). Whether this recruitment bottleneck was more severe in the LeConte population than in persistent enzootic MYL frog populations remains an important unanswered question.

Two results from the adult treatment experiments complicate the interpretation of the overall treatment effect. First, at the beginning of the experiment, frogs in the control group were captured and processed 1–3 days after frogs in the treated group. Because Bd loads in the population were increasing during this period, Bd loads on control frogs were somewhat higher than those on treated frogs. This could have exaggerated the subsequent differences in survival between control and treated frogs. Although we acknowledge this potential confounding effect, two factors suggest that the initial differences in Bd loads between control and treated frogs were not the primary cause of the lower survival of control frogs. First, pre-treatment Bd loads of control and treated frogs were very high, and given the relationship between Bd load and estimated survival in untreated control frogs (Figure 4c), survival of untreated frogs over the range of pre-treatment Bd loads observed in both groups would be expected to be near zero. Second, for the range of Bd load values that overlapped between frogs in the control and treated groups, treated frogs had much higher survival than control frogs. Both results suggest that the higher survival of treated frogs compared to control frogs during the post-treatment period was primarily due to the reduction in Bd loads caused by the treatment, and not the difference in pre-treatment Bd loads between control and treated frogs.

The second complicating result is the unexpectedly large number of non-experimental frogs captured during the post-treatment period. In untreated MYL frog populations, Bd epizootics typically result in the mortality of all, or nearly all, adults within one year (Vredenburg et al. 2010). Based on this, if the treatment increased frog survival, as predicted, then during the post-treatment period we would have captured primarily treated frogs, with control frogs and frogs that were not included in the experiment (i.e., untreated “non-experimental” frogs) being rare or absent. This outcome was observed in the upper basin, where during 2016-2018 we captured 81 treated, zero control, and eight non-experimental frogs. Although this same pattern was true in the lower basin for treated and control frogs (132 and 2 captured, respectively), we also captured 615 non-experimental frogs. Based on their sizes, most were older adults that had survived the epizootic, and the remainder were new recruits that had survived the epizootic as subadults or small adults. During the 2016-2018 period, frog-Bd dynamics in this non-experimental group were similar to those of the treated frogs, suggesting that these frogs had also mounted an effective adaptive immune response, and as a result, subsequently showed increased Bd resistance/tolerance and relatively high survival. The mechanism underlying the unexpectedly high survival of the non-experimental frogs during the 2015 Bd epizootic is unknown. In theory, treatment of a large fraction of the adult population could have reduced the pathogen pressure experienced by untreated frogs and increased their survival (Briggs et al. 2010). However, this would have increased the survival of control and non-experimental frogs, but only non-experimental frogs were captured in large numbers. In addition, despite similar treatments conducted in both the upper and lower basins, we observed a large number of non-experimental frogs only in the lower basin. Another possible cause of unexpectedly high survival of non-experimental frogs, and only in the lower basin, could be the higher habitat complexity that characterizes the lower basin. The upper basin contains a single lake and its associated inlet and outlet streams, but no adjacent ponds, meadows, or springs that provide suitable *R. sierrae* habitat. As a result, the entire frog population is restricted to the site at which the epizootic occurred. In contrast, the lower basin contains a diverse array of aquatic habitats, including two lakes, four ponds, and associated streams, marshes, and springs, all of which were used by *R. sierrae* prior to the epizootic. Although conjectural, it is possible that frogs in some of these associated habitats experienced lower pathogen pressure, lower Bd loads, and higher survival during the epizootic than frogs in the much larger lake-dwelling populations.

In summary, the two adult treatment experiments both altered frog-Bd dynamics in a predictable way, reducing Bd loads of treated versus control frogs and increasing frog survival. Unlike the short-term effects resulting from the treatment of early life stages, effects on adults persisted over the three-year post-treatment period, and were consistent with adults having mounted an effective adaptive immune response against Bd infection. However, both experiments failed in their ultimate objective to facilitate the long-term persistence of the study populations in an enzootic state. Despite evidence of successful reproduction in all post-treatment years, little recruitment of new adults occurred and few treated frogs remained after three years. Therefore, even when treatment increases adult survival, the high susceptibility of early life stages to chytridiomycosis (Bakar et al. 2016, Grogan et al. 2018b) will often limit recruitment and future population growth, precluding long-term population persistence.

The short-term effects of the Treasure treatment paralleled those of the LeConte treatment. Treatment substantially reduced Bd loads, and loads remained low one month following treatment. However, despite 48% of the treated frogs being recaptured one month after treatment, no treated frogs were detected during the subsequent two summers, indicating that in this population treatment did not increase longer-term survival. The relatively few frogs treated and the inability to conduct this treatment as an experiment with treated and control groups preclude strong conclusions, but the lack of frog survival one year after treatment indicates that the strong effects of treatment observed in the LeConte experiments are not universal, and may depend on the timing of the treatment relative to the onset of the epizootic (later in Treasure than LeConte) or the inherent susceptibility of the frog population to Bd infection (e.g., Savage and Zamudio 2011).

### Microbiome augmentation of subadult frogs

Results from the treatment experiments described above indicate that the effectiveness of treatments in changing long-term frog-Bd dynamics and facilitating population persistence depends heavily on the survival of subadult frogs and their recruitment into the adult population under post-epizootic conditions. Given the low immunocompetence of subadults against Bd (Rollins-Smith 1998, Grogan et al. 2018b), reducing Bd loads on subadults using itraconazole appears insufficient to keep loads low over the longer term and increase survival (see results of Barrett and Dusy treatments). The addition of protective probiotic bacteria to the frog skin microbiome may be a possible means to reduce susceptibility of this vulnerable life stage to chytridiomycosis and increase survival to adulthood (Harris et al. 2009, Bletz et al. 2013, Rebollar et al. 2020). In this application, the effectiveness of probiotics will depend critically on the ability by the added bacteria to establish on frog skin and maintain sufficiently high densities over the months or years of the subadult-to-adult transition.

The results from our microbiome augmentation experiment suggest that the addition of *J. lividum* to the frog skin microbiome following itraconazole treatment is insufficient to provide the long-term protection from Bd infection required to increase subadult survival. The relatively rapid decline of *J. lividum* concentrations on frogs in our study population suggests that the frog microbiome is resilient to changes in the species composition of symbiotic bacteria, and represents an important impediment to efforts to augment the microbiome with species that might confer increased protection from Bd (Küng et al. 2014). In addition, for several weeks immediately following probiotic exposure when *J. lividum* concentrations were relatively high, Bd loads on *J. lividum*-treated subadults increased quickly and those of control and wild subadults appeared unaffected. Therefore, the predicted protective effect of *J. lividum* on subadults was not realized. However, an unexpected outcome of the microbiome augmentation experiment was the rapid spread of *J. lividum* from exposed subadults to control and wild subadults. Control and wild frogs quickly developed *J. lividum* concentrations on their skin that were similar to those of frogs that were bathed in a concentrated *J. lividum* solution for several hours over a two-day period. Whether *J. lividum* was transferred via direct frog-to-frog contact or through the water is unknown. In conclusion, although the colonization of frogs by *J. lividum* did not appear to confer increased Bd resistance, its spread from *J. lividum*-exposed to unexposed frogs indicates that if a probiotic with long-term effectiveness against Bd infection is ever identified, its introduction into a frog population may be relatively straight-forward.

## Conclusions

Our experiments indicate that treatment of early life stage and adult MYL frogs with antifungal agents during or immediately following epizootics strongly altered frog-Bd dynamics over the short term. However, in the long term, all six treatments failed to allow treated populations to persist in the presence of Bd. Given this, recovery of MYL frogs will require other approaches that more effectively mitigate the impacts of Bd infection, in particular by increasing the survival and recruitment of early life stages. MYL frog recovery efforts conducted during the past 15 years indicate the potential of frog translocations using animals collected from populations that have rebounded during the two or more decades following past Bd epizootics (Knapp et al. 2016, Joseph and Knapp 2018). Frogs in these populations may have genotypes that are more resistant to or tolerant of Bd infection than those in Bd-naive populations (Knapp et al. 2016), providing an important advantage to translocated individuals. Use of such populations as sources of frogs for translocations is not universally effective in allowing population re-establishment (Joseph and Knapp 2018, see also Brannelly et al. 2016), but results from more than 20 translocations of adult MYL frogs conducted to date indicate a high probability of success. Encouragingly, these translocated populations are typically characterized by stable, enzootic frog-Bd dynamics, low-to-moderate Bd loads across a wide range of frog densities, and annual survival exceeding 50% (e.g., Joseph and Knapp 2018). Such translocations provide the best known opportunity to reestablish extirpated MYL frog populations across their historical range. Given possible evolution of increased resistance or tolerance in other amphibian species following exposure to Bd (Bataille et al. 2015, Savage and Zamudio 2016, Voyles et al. 2018), similar efforts might be applicable to the recovery of other Bd-endangered amphibian species (e.g., Brannelly et al. 2016, Mendelson III et al. 2019).

## Acknowledgements

The research described in this paper was supported by grants from Sequoia and Kings Canyon National Parks, National Science Foundation–National Institutes of Health Ecology of Infectious Disease program (EF-0723563), National Science Foundation Rapid Response Research program (IOS-1244804), and National Science Foundation Long-term Research in Environmental Biology program (DEB-1557190). Development of this paper was supported by Cooperative Agreement P19AC00789 from the National Park Service. The following people assisted with fieldwork: A. Adams, A. Beechan, D. Burkhart, K. Atkinson, I. Chellman, B. Currinder, C. Dorsey, M. Hernandez, B. Karin, N. Kauffman, A. Killion, J. Lester, A. Lindauer, S. Maple, M. Masten, D. Paolilli, W. Philbrook, G. Ruso, A. Stoerp, and L. Torres. L. Torres developed the *J. lividum* qPCR protocol while employed in the Vredenburg lab (San Francisco State University). M. Toothman in the Briggs lab (University of California-Santa Barbara) analyzed Bd swabs collected prior to 2016, and K. Rose and A. Barbella in the Knapp lab analyzed swabs collected thereafter. Research permits were provided by Sequoia and Kings Canyon National Parks, U.S. Fish and Wildlife Service, U.S. Forest Service, and the Institutional Animal Use and Care Committee at the University of California-Santa Barbara.

