## Supporting Information for "Effectiveness of antifungal treatments during chytridiomycosis epizootics in populations of an endangered frog"

### Author Contributions

Roland Knapp obtained the permits for all treatments, designed and implemented all of the treatments except at Treasure, conducted post-treatment surveys, managed the datasets, conducted final analyses of all of the treatment datasets except the capture-mark-recapture (CMR) dataset from the LeConte and Treasure treatments, and wrote most of the manuscript.

Maxwell Joseph assisted with the Barrett-2009 and Dusy-2010 treatments and associated post-treatment surveys, analyzed the CMR dataset from the LeConte experiment and wrote the associated text, provided input on most of the analyses, and reviewed all drafts of the manuscript.

Thomas Smith assisted with the LeConte and Treasure treatments, conducted post-treatment surveys for multiple treatments, analyzed the results of the Treasure treatment and wrote some of the associated text, conducted preliminary analyses of the Barrett-2009, Dusy-2010, and Dusy-2012 treatments, and reviewed all drafts of the manuscript.

Ericka Hegeman assisted with the Treasure treatment, conducted post-treatment surveys for multiple treatments, managed the datasets, conducted preliminary analyses of the Barrett-2009 and Dusy-2010 treatments and wrote some of the associated text, and reviewed all drafts of the manuscript.

Vance Vredenburg assisted with the design and implementation of the Dusy-2012 treatment, including supporting the development of the *J. lividum* qPCR assay, culturing of the *J. lividum* used in the treatment, and qPCR analysis of *J. lividum* samples.

James Erdman conducted pre- and post-treatment surveys at Treasure, implemented the Treasure treatment, and reviewed all drafts of the manuscript.

Daniel Boiano assisted with the LeConte treatment, facilitated the National Park Service permitting of all treatments conducted in Sequoia and Kings Canyon National Parks, and reviewed all drafts of the manuscript.

Andrea Jani assisted with the LeConte treatment, and conducted the Dusy Basin zoospore pool study, provided the associated dataset, conducted preliminary analyses, and wrote some of the associated text, and reviewed all drafts of the manuscript.

Cheryl Briggs provided modeling results that motivated the treatments, assisted with the design of treatments, and reviewed all drafts of the manuscript.

### Methods

#### Estimating the zoospore pool from water samples

We sampled the zoospore pool in each of the five ponds included in the Dusy itraconazole treatment experiment (Table 1) before and after the treatments (July 23-25 and August

21-24, respectively). Water samples (six per pond) were collected by filtering pond water through a 0.22 mm pore polyethersulfone filter (Sterivex-GP; Millipore), until the filter clogged. Filters were immediately amended with sucrose lysis buffer (40 mmol/l 1 EDTA, 50 mmol/l 1 Tris-HCl, 750 mmol/l sucrose, pH adjusted to 8.0). We extracted DNA using the DNEasy Blood and Tissue kit (Qiagen). Bd concentrations in water samples (environmental "Bd loads") were quantified using qPCR (see Jani et al. 2017 for details), and Bd load was normalized to a 1-liter sample volume. For each sample, we ran three technical replicates, each from an independent sample dilution and on an independent PCR run. To minimize PCR inhibition, we diluted DNA extracts 50-fold based on pilot tests using undiluted, ten-, fifty-, and one hundred-fold dilutions. Finally, we included 21 negative controls: nine no-template-controls (three per PCR plate) and 12 field-collected negative controls (six water samples collected from a pond where all frogs were Bd-negative, each with 2 technical replicates).

The 21 negative controls yielded no false-positive PCR reactions. However, it is common for technical replicates of environmental DNA samples to have a high rate of false-negatives due to low quantities of target DNA or PCR inhibitors in samples (Mosher et al. 2018), and this was the case in our study. Despite the fact that all five study ponds contained relatively large numbers of early life stage *R. sierrae* characterized by high Bd loads, of the 180 total replicates (5 ponds x 6 water samples x 3 technical replicates x 2 time periods (before and after treatment)), 49% had a Bd load = 0. Because the five ponds in the experiment were clearly Bd-positive, we considered replicates with Bd load = 0 to be false negative replicates, which we excluded from the analysis of the effect of itraconazole treatment on the Bd zoospore pool. This resulted in one pond assigned to the treated group being dropped from the analysis due to a lack of any Bd-positive replicates. The effect of itraconazole treatment on Bd concentrations on filters was evaluated using the model *bd_load ~ (pre_post x treatment) + (1 | sample_id)* (family = negative binomial, pre_post = [before treatment, after treatment], treatment

= [treated, control], sample_id included as a group-level effect to account for technical replicates).

#### *Janthinobacterium lividum* qPCR assay

The *J. lividum* used to develop the real-time PCR assay was a strain provided by Reid Harris, Department of Biology, James Madison University. *J. lividum* DNA was extracted from a pure culture of this strain using a Qiagen DNeasy blood and tissue kit following the manufacturer's protocol. The DNA collected was amplified using methods in Harris et al. (2009). The PCR product was viewed on a 1% agarose gel, and the 500 bp product was sequenced.

Primers were designed to be specific to *J. lividum* and sequences were:

- Jliv_For3 ATGCCACCGACGGCTACCA
- Jliv_Rev1 ACGGCGGGATGGTCATCAC

The minor groove binder probe sequence was:

- JLIVT 6FAM AACATCGTTTGCTGTCCGTTGA MGBNFQ

After assay optimization, the 25 $\mu$L reaction volumes included 0.5 $\mu$L of each primer at a concentration of 10 $\mu$M, 0.375 $\mu$L of MGB probe at a concentration of 250 nM, 12.5 $\mu$L of Taqman MasterMix (Applied Biosystems), 1.25 $\mu$L BSA, and 5 $\mu$L of template. The amplification conditions consisted of an initial cycle of 2 min at 50°C and 10 min at 95°C, followed by 50 cycles at 95°C for 15 s, 58°C for 30 s, and 65°C for 45 s. To create standards, DNA was extracted from pure cultures of *J. lividum* with an UltraClean microbial DNA isolation kit (MoBio), and diluted to $10^4$, $10^3$, $10^2$, and $10^1$ *J. lividum* genome equivalents. A standard curve was generated for each 96-well plate to estimate the number of *J. lividum* genome equivalents in sample extracts.

#### Hidden Markov model for the LeConte treatment experiment

We tracked the fates of individual animals in the LeConte population using a multi-state hidden Markov model. We consider three possible observations for each individual $i = 1, ..., M$ in primary period $t = 1, ..., T$, on secondary period $j = 1, ..., J_t$, where $T$ is the total number of primary periods and $J_t$ is the number of secondary periods in primary period $t$:

- $y_{i,t,j} = 1$ detected at the upper site
- $y_{i,t,j} = 2$ detected at the lower site
- $y_{i,t,j} = 3$ not detected

We use parameter-expanded data augmentation to account for the fact that the total number of adults in the population is unknown (Royle and Dorazio 2012). Across the entire time period of the study, we assume $N_s$ unique individuals have been alive at either site. We observe $N$ unique individuals across all surveys, where $N \leq N_s$. An estimate of $N_s$ can be acquired by augmenting the observed capture histories with additional capture histories that consist entirely of non-detections, thus modeling a large number $M > N_s$ of individuals, $M - N_s$ of which never existed (Royle 2009). Here, $M$ was chosen to be 2182 (1212 observed unique individuals plus 970 augmented individuals). We verified that posterior estimates of $N_s$ were much less than $M$ to avoid problems on the boundary of this augmented parameter space (Dennis et al. 2015).

We denote the true state of individual $i$ in primary period $t$ as $s_{i,t}$. We assume that within a primary period, the state of each individual does not change. This assumption is justified by the short time intervals between secondary periods within primary periods. Four states are possible:

- $s_{i,t} = 1$ alive at the upper site

- $s_{i,t} = 2$ alive at the lower site
- $s_{i,t} = 3$ not recruited
- $s_{i,t} = 4$ dead

**Observation model**

An emission matrix $\mathbf{\Omega}^{(t)}$ links observations to hidden states for primary period $t$. The rows in $\mathbf{\Omega}^{(t)}$ correspond to the state of an individual in primary period $t$, and the columns correspond to observation probabilities such that the entry in the $m^{th}$ row, $n^{th}$ column is $Pr(y_{i,t,j} = n \mid s_{i,t} = m)$:

$$\mathbf{\Omega}^{(t)} = \begin{array}{c} \\ \end{array}\begin{pmatrix} \text{Detected: upper} & \text{Detected: lower} & \text{Not detected} \\ p_t & 0 & 1 - p_t \\ 0 & p_t & 1 - p_t \\ 0 & 0 & 1 \\ 0 & 0 & 1 \end{pmatrix} \begin{array}{l} \\ \text{Alive: upper} \\ \text{Alive: lower} \\ \text{Not recruited} \\ \text{Dead} \end{array}$$

Here, $p_t$ is the probability of detection for an individual if it is alive in primary period $t$. We allowed detection probabilities to vary over time (Joseph and Knapp 2018):

$$\text{logit}(p_t) = \alpha_0 + \epsilon_t^{(p)},$$

where $\alpha_0$ is an intercept parameter, and $\epsilon_t^{(p)}$ is an adjustment on detection probability for primary period $t$.

#### State model

The hidden states of each individual evolve as a Markov process with transition matrix $\mathbf{\Psi}^{(i,t)}$, where the element in the $m^{th}$ row, $n^{th}$ column is $\Pr(s_{i,t+1} = n \mid s_{i,t} = m)$:

$$\begin{array}{c c} & \begin{array}{cccc} \text{Alive: upper} & \text{Alive: lower} & \text{Not recruited} & \text{Dead} \end{array} \\ \mathbf{\Psi}^{(i,t)} = & \begin{pmatrix} \phi_{i,t}(1-\nu^{(l)}) & \phi_{i,t}\nu^{(l)} & 0 & 1-\phi_{i,t} \\ \phi_{i,t}\nu^{(u)} & \phi_{i,t}(1-\nu^{(u)}) & 0 & 1-\phi_{i,t} \\ \gamma_t\rho^{(u)} & \gamma_t(1-\rho^{(u)}) & 1-\gamma_t & 0 \\ 0 & 0 & 0 & 1 \end{pmatrix} \begin{array}{c} \text{Alive: upper} \\ \text{Alive: lower} \\ \text{Not recruited} \\ \text{Dead} \end{array} \end{array}$$

Here, $\phi_{i,t}$ is the probability of survival, $\nu^{(l)}$ and $\nu^{(u)}$ are the probabilities of moving to the lower or upper site respectively (conditional on survival), $\gamma_t$ is the probability of recruitment, and $\rho^{(u)}$ is the probability of recruiting into the upper site conditional on recruitment.

Survival probabilities were modeled as a function of Bd load:

$$\text{logit}(\phi_{i,t}) = \beta_0 + \beta^{(g)}_{\text{g[i]}} + \beta^{(z)}_{\text{g[i]}} z_{i,t},$$

where $\beta_0$ is an intercept parameter, $\beta^{(g)}_{\text{g[i]}}$ is an adjustment for group $g$, $g[i]$ is the group that individual $i$ belongs to, $\beta^{(z)}_{\text{g[i]}}$ is the effect of Bd load on group $g$, and $z_{i,t}$ is the Bd load of individual $i$ in primary period $t$.

We treat the field experiment in 2015 as the first primary period, for which states are known for experimental animals. For example, if individual $i$ was captured and released at the upper site, then we know that $s_{i,t} = 1$. Initial states are not known for non-experimental animals, which could have been alive at either site (states 1 or 2), or in the not recruited class (state 3). Note that we are only interested in modeling the state and capture histories of animals that might have been alive. We assigned a Dirichlet(1, 1, 1) prior for the initial state distribution for non-experimental frogs, which assigns equal prior density to each initial state.

#### Bd load model

We modeled Bd loads as being normally distributed on a transformed scale. Raw Bd loads were transformed using a $\log_{10} + 1$ transformation, then centered and scaled to have mean zero and unit standard deviation (in an attempt to avoid ill-conditioning, as expected Bd load is used in the detection and survival model components). Let $z_{i,t}^{\text{obs}}$ represent the observed transformed Bd load, and $z_{i,t}$ represent the expected value for individual $i$ on primary period $t$. The observation model for Bd loads of detected individuals was Gaussian on the transformed scale:

$$z_{i,t}^{\text{obs}} \sim \text{Normal}(z_{i,t}, \sigma),$$

where $\sigma$ is an observation-level standard deviation parameter.

Expected Bd load was modeled as a function of treatment, primary period, and individual identity:

$$z_{i,t} = \mu_t + \beta_{\text{trt}[i]} + \epsilon_i,$$

where $\mu_t$ is a time-varying intercept, $\beta_{\text{trt}[i]}$ is an adjustment for the treatment group of individual $i$ (denoted trt$[i]$) to account for differences in mean Bd loads between treated, control, and non-experimental animals, and $\epsilon_i$ is an individual-level adjustment.

##### Prior distributions

We expected movement among sites to be rare, so both movement parameters ($\nu^{(l)}$ and $\nu^{(u)}$) were assigned Beta(2, 20) priors. Primary period and individual-level adjustments were modeled using zero-mean normal distributions with unknown standard deviations specific to the process of interest, e.g., for the probability of entering the population: $[\lambda_{1:T}] = \prod_{t=1}^{T} \text{Normal}(\lambda_t | 0, \sigma^{(\lambda)})$. Standard deviation parameters were given unit scale half normal priors, and all remaining parameters were given unit scale normal priors.

##### Inference

We sampled from the posterior distribution of this model using dynamic Hamiltonian Monte Carlo in Stan. We drew 3000 iterations for each of four chains, using a maximum treedepth of 11 and an adapt_delta value of 0.99. Convergence was checked by visual inspection of traceplots and with Rhat values, using Rhat $< 1.1$ as a threshold. Models were fit using the rstan R package, version 2.21.2 (Stan Development Team 2020).

### Results

#### Estimating the zoospore pool from water samples

Before treatment, zoospore pools (measured as Bd load on collected filters) in the ponds assigned to the control and treated groups were similar (Figure S2). After treatment, zoospore pools in control ponds may have increased slightly, but remained relatively constant in treated ponds (Figure S2). Model results indicated that the estimated effects of treatment, basin, and the (treatment x basin) interaction term were all unimportant (Table S5). Therefore, assuming that the sampling method was adequate to accurately quantify pond-wide zoospore concentrations, the treatment of even a relatively large fraction of the resident *R. sierrae* in the study ponds did not measurably alter the zoospore pools. To avoid the high number of false-negative filters obtained using our methods, future studies attempting to quantify zoospore pools should consider using methods that allow filtering of larger volumes of water.

Stan Development Team. 2020. RStan: The R interface to Stan.

### Tables

#### Table S1

Characteristics of all sites used in the antifungal treatment experiments

| Basin | Site ID | Experiment | Life stage | Category | Elevation | Depth | Area |
|---|---|---|---|---|---|---|---|
| Barrett | 11469 | itraconazole | early | treated | 3383 | 2.7 | 3875 |
| Barrett | 11491 | itraconazole | early | treated | 3530 | 5.0 | 2316 |
| Barrett | 11493 | itraconazole | early | treated | 3459 | 0.6 | 269 |
| Barrett | 11470 | itraconazole | early | control | 3383 | 5.0 | 3998 |
| Barrett | 10222 | itraconazole | early | control | 3554 | 5.2 | 10568 |
| Barrett | 11495 | itraconazole | early | control | 3459 | 1.0 | 970 |
| Dusy | 11518 | itraconazole | early | treated | 3408 | 1.0 | 2002 |
| Dusy | 11526 | itraconazole | early | treated | 3219 | 1.9 | 2966 |
| Dusy | 11506 | itraconazole | early | treated | 3469 | 1.8 | 1414 |
| Dusy | 11517 | itraconazole | early | control | 3395 | 0.8 | 816 |
| Dusy | 11525 | itraconazole | early | control | 3158 | 1.2 | 2604 |
| LeConte | 10101 | itraconazole | adult | treated | 3213 | 1.5 | 5187 |
| LeConte | 10100 | itraconazole | adult | treated | 3298 | 14.9 | 25974 |
| Treasure | 50839 | itraconazole | adult | treated | 3410 | 11.0 | 34317 |
| Dusy | 11518 | itracon + Jliv | subadult | treated | 3408 | 1.0 | 2002 |

Life stage = "early" indicates tadpoles and subadults

Treatment = "itracon + Jliv" indicates the use of itraconazole and *J. lividum*

Units for Elevation and Depth are meters and those for Area are $m^2$

### Table S2

Timelines for antifungal treatment experiments conducted in the lower and upper basins of the LeConte study area in 2015.

(a) Lower basin

| **Day 1 (24-Aug)** | **Day 2 (25-Aug)** | **Day 3 (26-Aug)** | **Day 4 (27-Aug)** | **Day 5-7 (28-30 Aug)** | **Day 8 (31-Aug)** | **Day 9 (1-Sep)** |
|---|---|---|---|---|---|---|
| Captured 50 frogs for "treated" group, Day 1 frogs swabbed and treated. | Captured 157 frogs for "treated" group, Day 2 frogs swabbed, Day 1 & 2 frogs treated. | Captured 152 frogs for "treated" group, Day 3 frogs swabbed, Day 1-3 frogs treated. | Captured 102 frogs for "control" group, swabbed and released. Day 1-3 frogs treated. | Day 1-3 frogs treated. | Day 1-3 frogs treated. Swabbed subset of Day 1, 2, and 3 frogs. | Day 1-3 frogs treated, released back into lakes. |

(b) Upper basin

| **Day 1 (8-Sep)** | **Day 2 (9-Sep)** | **Day 3 (10-Sep)** | **Day 4-6 (11-13 Sep)** | **Day 7 (14-Sep)** | **Day 8 (15-Sep)** |
|---|---|---|---|---|---|
| Captured 45 frogs for "treated" group, Day 1 frogs swabbed and treated. | Captured 161 frogs for "treated" group, Day 2 frogs swabbed, Day 1 & 2 frogs treated. | Captured 74 frogs for "control" group, swabbed and released, Day 1 & 2 frogs treated. | Day 1 & 2 frogs treated. | Day 1 & 2 frogs treated. Swabbed subset of Day 1 & 2 frogs. | Day 1 & 2 frogs treated, released back into lakes. |

#### Table S3

For the itraconazole treatment experiments in Barrett and Dusy basins, results of model comparing Bd loads on frogs in ponds assigned to control and treated groups immediately before the treatment period (model family is negative binomial).

| | Estimate | Est. Error | lo95%CI | up95%CI | Rhat | Bulk ESS | Tail Ess |
|---|---|---|---|---|---|---|---|
| **Population-level effects** | | | | | | | |
| Intercept | 11.99 | 0.55 | 11.05 | 13.21 | 1.00 | 2273 | 1948 |
| treatment(treated) | 0.15 | 0.77 | -1.39 | 1.64 | 1.00 | 1956 | 1993 |
| basin(dusy) | -1.36 | 0.70 | -2.81 | -0.02 | 1.00 | 2058 | 1797 |
| treatment(treated):basin(dusy) | 1.36 | 0.93 | -0.54 | 3.17 | 1.00 | 1842 | 1705 |
| **Family-specific parameters** | | | | | | | |
| overdispersion | 0.30 | 0.04 | 0.23 | 0.39 | 1.00 | 2777 | 2617 |

### Table S4

For the itraconazole treatment experiments in Barrett and Dusy basins, results of model comparing Bd loads on frogs assigned to the treated group from before versus the end of the treatment period (model family is negative binomial).

| | Estimate | Est. Error | lo95%CI | up95%CI | Rhat | Bulk ESS | Tail Ess |
|---|---|---|---|---|---|---|---|
| **Population-level effects** | | | | | | | |
| Intercept | 13.46 | 0.27 | 12.96 | 14.00 | 1.00 | 3780 | 3096 |
| overdispersion-Intercept | -1.59 | 0.12 | -1.84 | -1.35 | 1.00 | 4561 | 3194 |
| stage(tadpole) | -2.28 | 0.29 | -2.87 | -1.74 | 1.00 | 3248 | 3010 |
| trt_period(end) | -2.56 | 0.22 | -2.98 | -2.11 | 1.00 | 3179 | 2841 |
| basin(dusy) | 1.71 | 0.19 | 1.35 | 2.06 | 1.00 | 3680 | 3357 |
| trt_period(end):basin(dusy) | -4.82 | 0.45 | -5.68 | -3.88 | 1.00 | 3125 | 2537 |
| overdispersion-stage(tadpole) | 1.36 | 0.14 | 1.10 | 1.64 | 1.00 | 4018 | 2923 |
| overdispersion-trt_period(end) | -1.11 | 0.11 | -1.34 | -0.90 | 1.00 | 3478 | 3082 |
| overdispersion-basin(dusy) | -0.83 | 0.12 | -1.05 | -0.59 | 1.00 | 4047 | 2579 |

### Table S5

For the Dusy Basin itraconazole treatment experiment, results of model comparing zoospore pools of ponds assigned to the control and treated groups before and after treatment (model family is negative binomial).

| | Estimate | Est. Error | lo95%CI | up95%CI | Rhat | Bulk ESS | Tail Ess |
|---|---|---|---|---|---|---|---|
| **Group-level effects** | | | | | | | |
| sd(Intercept) | 1.98 | 0.28 | 1.46 | 2.55 | 1.01 | 471 | 618 |
| **Population-level effects** | | | | | | | |
| Intercept | 8.65 | 0.64 | 7.42 | 9.89 | 1.00 | 558 | 987 |
| pre_post(post) | 1.74 | 0.94 | -0.13 | 3.54 | 1.01 | 467 | 727 |
| tmt(treated) | 1.19 | 1.02 | -0.87 | 3.20 | 1.01 | 627 | 886 |
| pre_post(post):tmt(treated) | -2.54 | 1.28 | -5.07 | 0.00 | 1.01 | 525 | 947 |
| **Family-specific parameters** | | | | | | | |
| overdispersion | 3.04 | 0.64 | 1.87 | 4.42 | 1.00 | 787 | 703 |

#### Table S6

For the LeConte Basin itraconazole treatment experiment, results of model comparing Bd loads on frogs assigned to the control and treated categories immediately before the treatment period (model family is negative binomial).

| | Estimate | Est. Error | lo95%CI | up95%CI | Rhat | Bulk ESS | Tail Ess |
|---|---|---|---|---|---|---|---|
| **Population-level effects** | | | | | | | |
| Intercept | 17.01 | 0.13 | 16.75 | 17.27 | 1.00 | 2262 | 2396 |
| location(upper) | 0.03 | 0.20 | -0.36 | 0.44 | 1.00 | 1663 | 2309 |
| group(treated) | -0.90 | 0.16 | -1.21 | -0.60 | 1.00 | 2185 | 2317 |
| location(upper):group(treated) | 0.51 | 0.25 | 0.00 | 1.01 | 1.00 | 1679 | 2222 |
| **Family-specific parameters** | | | | | | | |
| overdispersion | 0.54 | 0.03 | 0.49 | 0.59 | 1.00 | 3333 | 2393 |

#### Table S7

For the LeConte Basin itraconazole treatment experiment, results of model comparing Bd loads on frogs assigned to the treated group from before versus the end of the treatment period (model family is negative binomial).

| | Estimate | Est. Error | lo95%CI | up95%CI | Rhat | Bulk ESS | Tail Ess |
|---|---|---|---|---|---|---|---|
| **Population-level effects** | | | | | | | |
| Intercept | 16.11 | 0.11 | 15.90 | 16.32 | 1.00 | 4204 | 3258 |
| location(upper) | 0.54 | 0.19 | 0.16 | 0.93 | 1.00 | 3486 | 3216 |
| trt_period(endtreat) | -2.66 | 0.22 | -3.07 | -2.23 | 1.00 | 3170 | 3162 |
| location(upper):trt_period(endtreat) | 1.15 | 0.38 | 0.41 | 1.89 | 1.00 | 3040 | 2980 |
| **Family-specific parameters** | | | | | | | |
| overdispersion | 0.31 | 0.01 | 0.29 | 0.34 | 1.00 | 3988 | 2862 |

### Table S8

For the LeConte Basin itraconazole treatment experiment, results of model comparing Bd loads on frogs in the treated group that survived versus died (model family is bernoulli).

| | Estimate | Est. Error | lo95%CI | up95%CI | Rhat | Bulk ESS | Tail Ess |
|---|---|---|---|---|---|---|---|
| **Population-level effects** | | | | | | | |
| Intercept | -1.90 | 0.90 | -3.70 | -0.20 | 1.00 | 2120 | 2090 |
| lbd_load | 0.09 | 0.14 | -0.18 | 0.36 | 1.00 | 2050 | 2081 |
| location(upper) | 0.11 | 2.02 | -3.85 | 4.15 | 1.00 | 1460 | 1663 |
| lbd_load:location(upper) | 0.11 | 0.29 | -0.48 | 0.68 | 1.00 | 1434 | 1693 |

### Table S9

For the Treasure Lakes Basin itraconazole treatment, results of model comparing Bd loads on frogs before versus the end of the treatment period (model family is negative binomial).

| | Estimate | Est. Error | lo95%CI | up95%CI | Rhat | Bulk ESS | Tail Ess |
|---|---|---|---|---|---|---|---|
| **Population-level effects** | | | | | | | |
| Intercept | 15.45 | 0.22 | 15.05 | 15.91 | 1.00 | 3452 | 2500 |
| trt_period(after) | -1.35 | 0.41 | -2.10 | -0.50 | 1.00 | 3817 | 2448 |
| **Family-specific parameters** | | | | | | | |
| overdispersion | 0.29 | 0.03 | 0.23 | 0.35 | 1.00 | 3580 | 2946 |

### Table S10

For the Treasure Lakes Basin itraconazole treatment, results of model evaluating predictors of treatment effectiveness (model family is gaussian).

| | Estimate | Est. Error | lo95%CI | up95%CI | Rhat | Bulk ESS | Tail Ess |
|---|---|---|---|---|---|---|---|
| **Population-level effects** | | | | | | | |
| Intercept | 3.13 | 2.68 | -2.15 | 8.40 | 1.00 | 1800 | 2078 |
| capture_bdload_std | -5.93 | 4.88 | -15.25 | 4.22 | 1.00 | 1357 | 1773 |
| days_inside | -1.42 | 0.43 | -2.26 | -0.57 | 1.00 | 1820 | 2119 |
| capture_bdload_std:days_inside | 0.56 | 0.75 | -0.99 | 1.99 | 1.00 | 1351 | 1549 |
| **Family-specific parameters** | | | | | | | |
| sigma | 3.10 | 0.43 | 2.40 | 4.07 | 1.00 | 2484 | 2221 |

### Table S11

For the Dusy Basin microbiome augmentation experiment, results of model comparing Bd loads on frogs assigned to the control and treated groups immediately before the itraconazole treatment period (model family is negative binomial).

| | Estimate | Est. Error | lo95%CI | up95%CI | Rhat | Bulk ESS | Tail Ess |
|---|---|---|---|---|---|---|---|
| **Population-level effects** | | | | | | | |
| Intercept | 13.97 | 0.22 | 13.57 | 14.42 | 1.00 | 3514 | 2409 |
| expt_trt(treated) | 0.17 | 0.27 | -0.37 | 0.68 | 1.00 | 3111 | 2460 |
| **Family-specific parameters** | | | | | | | |
| overdispersion | 0.82 | 0.12 | 0.61 | 1.07 | 1.00 | 3564 | 2992 |

#### Table S12

For the Dusy Basin microbiome augmentation experiment, results of model comparing Bd loads on frogs assigned to the treated category from immediately before versus at the end of the itraconazole treatment (model family is zero-inflated negative binomial).

| | Estimate | Est. Error | lo95%CI | up95%CI | Rhat | Bulk ESS | Tail Ess |
|---|---|---|---|---|---|---|---|
| **Population-level effects** | | | | | | | |
| Intercept | 14.15 | 0.13 | 13.90 | 14.42 | 1.00 | 5206 | 2934 |
| days(0) | -8.92 | 0.22 | -9.35 | -8.47 | 1.00 | 3798 | 2834 |
| **Family-specific parameters** | | | | | | | |
| overdispersion | 1.10 | 0.16 | 0.79 | 1.43 | 1.00 | 3980 | 2962 |
| zi | 0.21 | 0.04 | 0.13 | 0.29 | 1.00 | 3980 | 2480 |

### Figures

#### Figure S1

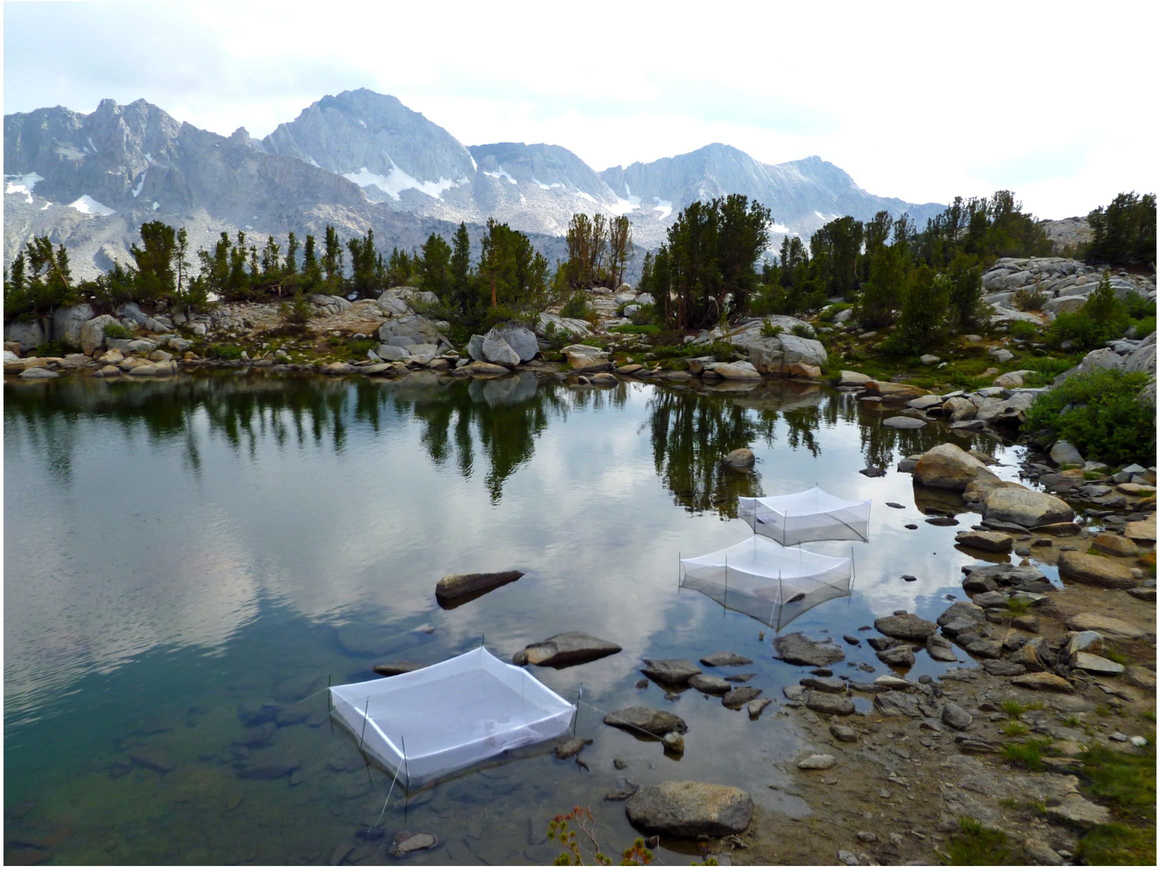

Photograph showing mesh pens used to hold subadult *R. sierrae* during the 2012 microbiome augmentation experiment in Dusy Basin.

#### Figure S2

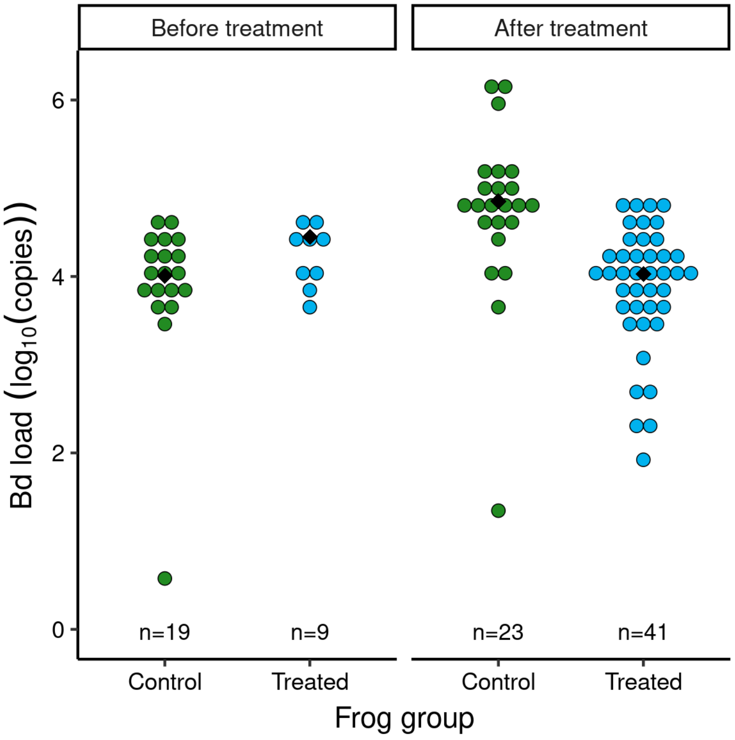


In the Dusy Basin study ponds assigned to the control or treated groups, zoospore pools measured before and after itraconazole treatment of *R. sierrae.* The y-axis displays Bd load per water sample, normalized to a 1-liter sample volume. Each dot represents a single sample, and median values for each treatment period are indicated with a black diamond. The number of samples included is displayed above the x-axis.

#### Figure S3

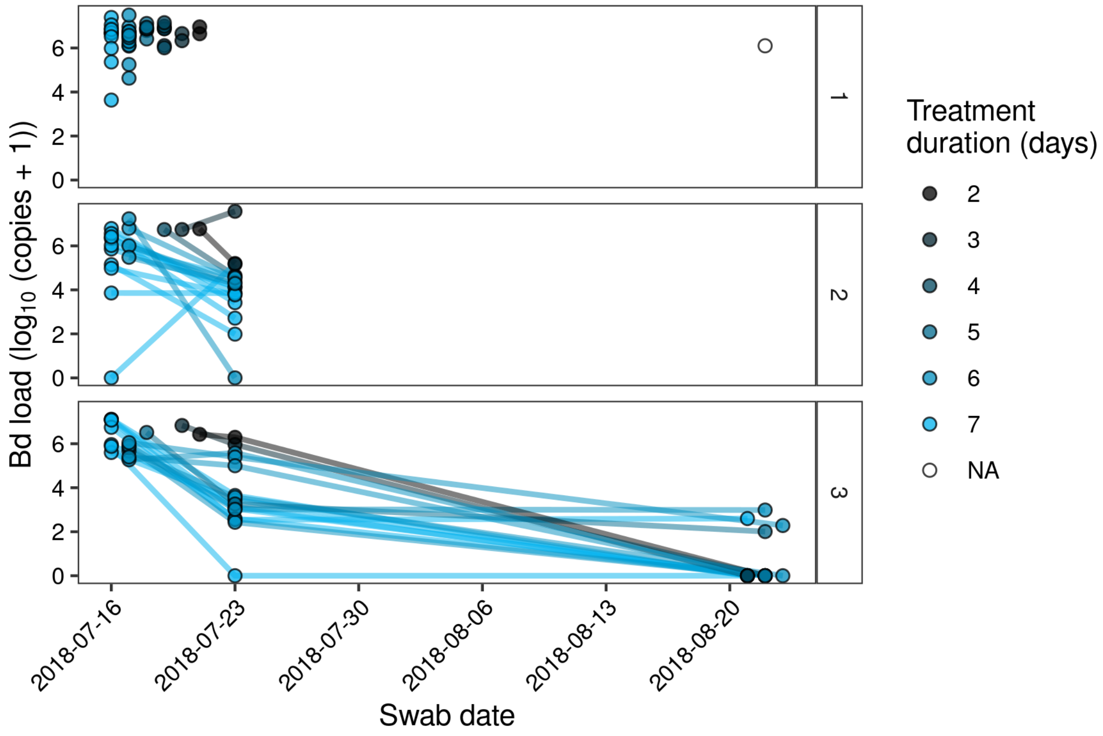


For all adult *R. sierrae* in the 2018 Treasure Lakes itraconazole treatment, Bd loads over a two month period that includes the July treatment (16 July to 23 July 2018) and August follow-up surveys. Points from the same frog are connected by a line. Panel labels are as follows: 1 = frog that died during the treatment ("non-survivor"), 2 = survivor that was not recaptured during the post-release survey in August, and 3 = survivor that was recaptured during the post-release survey. The single non-experimental frog captured in August was not included in the treatment (treatment duration = "NA") and is included in panel 1.
